# Make or break: The influence of expected challenges and rewards on the motivation and experience associated with cognitive effort exertion

**DOI:** 10.1101/2023.12.05.570154

**Authors:** Yue Zhang, Xiamin Leng, Amitai Shenhav

## Abstract

Challenging goals can induce harder work but also greater stress, in turn potentially undermining goal achievement. We sought to examine how mental effort and subjective experiences thereof interact as a function of challenge level and the size of the incentives at stake. Participants performed a task that rewarded individual units of effort investment (correctly performed Stroop trials) but only if they met a threshold number of correct trials within a fixed time interval (challenge level). We varied this challenge level (Study 1, N = 40), and the rewards at stake (Study 2, N = 79), and measured variability in task performance and self-reported affect across task intervals. Greater challenge and higher rewards facilitated greater effort investment but also induced greater stress, while higher rewards (and lower challenge) simultaneously induced greater positive affect. Within intervals, we observed an initial speed up then slowdown in performance, which could reflect dynamic reconfiguration of control. Collectively, these findings further our understanding of the influence of task demands and incentives on mental effort exertion and wellbeing.

---

Goal attainment in everyday life requires exerting cognitive control. We exert control to fuel the necessary performance to reach the goal. However, control is effortful and how much control a person is willing to invest in reaching a goal depends on how motivated they are (Botvinick & Braver, 2015; Shenhav et al., 2017). Recent work has characterized the motivational factors that determine effort investment in a control-demanding task (e.g., Stroop-like tasks that require an individual to respond to a target feature while ignoring a distractor) (Botvinick & Braver, 2015). This work has shown that people adjust their level of control allocation based on performance-outcome contingencies (e.g., monetary rewards and penalties) and the expected difficulty of the task (e.g., target-distractor incongruency) in order to maximize performance towards a given goal (Bugg et al., 2011; Krebs et al., 2010; Leng et al., 2021). However, how control allocation adjusts to the level of challenge^1^ presented by the goal itself (i.e., how much effort is required to meet the goal) remains unclear. The current study manipulated challenge and incentive level to investigate influence on moment-to-moment exertion of mental effort, as well as affective experiences, in a novel experimental paradigm.

A large body of work in the field of organizational behavior shows that setting a challenging but attainable goal leads to better performance, relative to having no goal, a goal that is too challenging (e.g., a hard-to-meet performance target), or a goal that is not challenging enough (e.g., an easy-to-meet performance target) (Locke, 1968). Although a substantial body of work has gone into establishing and testing the goal setting theory, there are important elements of this theoretical framework that remain underexplored, in part due to limitations of past experiments. For instance, research on goal-setting often examines goal commitment over long timescales and were almost always based on self-report while ignoring behavioral indicators, such as how much and how long effort was exerted (Klein et al., 2013). It is therefore largely unknown how variability in challenge levels translate into within-participant changes in trial-by-trial performance on a control-demanding task. Recent approaches to studying interactions between motivation and cognitive control offer this additional level of granularity (e.g., by revealing how accuracy and response time vary as a function of incentive level; see Botvinick & Braver, 2015), but have yet to examine this critical component of motivation’s role in shaping control, nor how it interacts with the incentives for performance.

Several lines of work offer clues as to how expected challenges and incentives will interact to determine control allocation. Goal-setting theories, as well as an analogous line of research under the framework of Motivational Intensity Theory (Richter et al., 2016), predict that increasing levels of expected incentives and expected challenge should promote greater investment of mental effort, up to the point where it is no longer efficacious for improving performance (i.e., when the goal is increasingly impossible to meet), after which these theories predict that control will be divested (Shenhav et al., 2021; Silvestrini et al., 2023). This prediction has yet to be tested directly in the context of traditional cognitive control tasks. It is therefore also unclear to what extent increasingly challenging goals lead people to adjust the *type* of control they invest (e.g., their focus on speed versus accuracy) and how they dynamically adjust these control levels as they approach and after they have met their goal.

Separate from their role in motivating effort, it is also known that challenging tasks can induce negative affective experiences such as feelings of stress, which could in turn serve to undermine performance (Byron et al., 2018; Espedido & Searle, 2018). The factors that determine such feelings of acute stress are poorly understood, in part because these experiences are either not measured (as in the majority of research on motivation-control interactions) or are measured at a wide temporal scale (e.g., at the level of an experiment; Harvey & Victoravich, 2009; Henkel & Hinsz, 2004). It thus remains to be determined to what extent challenge level (and ensuing performance, including whether one succeeds or fails at meeting their goal) influence task-related experiences of both positive and negative affect. A particularly intriguing and as-yet-unaddressed question relates to how monetary incentives and challenge level might interact to determine momentary affective experiences of a task. Higher monetary reward should amplify one’s achievement if a goal is attained, leading to greater positive affect, but how these incentive levels affect stress is less clear. Past work suggests two intuitive but diametrically opposed hypotheses: greater reward incentives could increase the amount of stress experienced when viewed as higher stakes (similar to high stakes testing or competition conditions; Heissel et al., 2021; Yu, 2015). However, individual differences exist (Heissel et al., 2021) and reward incentives could also decrease the level of stress.

Here, we sought to build on past work to examine how performance and affect vary, and potentially interact, within a given person based on the level of challenge and stakes they are facing in performing a cognitively demanding task. Across two studies, we examine the influence of expected challenge level on (1) how much effort a person exerts and (2) the subjective affective experiences they feel while doing so, focusing on acute stress. We devised a timed, incentivized cognitive control task, wherein participants had to meet a specific goal threshold (number of correct trials) to receive their accumulated rewards. Throughout the experiment, we varied this goal threshold (i.e., challenge level) and measured how this influenced performance within and across time intervals, and how it influenced self-reported affective experiences. In Experiment 2, we additionally varied the amount of reward received for each correct response, to test how levels of expected reward interacts with challenge level in impacting effort and affect.

Consistent with past research on goal setting (Locke & Latham, 2019), we observed increased motivation with more challenging goals, such that participants would put in more effort and persist longer with their effort to achieve a higher goal. We also found better performance in high reward conditions. At the same time, we found that participants felt more stressed with both higher challenge level and higher monetary stakes, but that these factors diverged in their influences on positive affect, with greater challenge producing less positive affect and greater reward producing more. We further exploited our unique experimental approach to examine how goal proximity influences task performance, finding that participants initially sped up during the task then slowed down as they approached the goal, potentially consistent with adjustments in response threshold and evidence accumulation rate. In addition, we were also able to evaluate how task performance interacts with challenge level to influence affective experiences. Collectively, this work shows the impact of task demands and incentives on trial-to-trial exertion of mental effort and affective experiences through a novel experimental paradigm.

## Experiment 1: The effect of challenge level on performance and affective experiences

### Methods

#### Participants

We recruited 40 participants online through Prolific (https://www.prolific.co/). The sample size was determined based on prior studies that have established effects of stakes on control using a similar design (Frömer et al., 2021; Leng et al., 2021). All participants indicated that they had normal or corrected-to-normal vision and color vision prior to completing the study. Consent was given in compliance with Brown University’s Institutional Review Board (1606001539). Participants were compensated for their time and received an additional monetary bonus based on their performance of the task. Two participants were excluded from our analyses because their affective ratings were uniform at the default rating (5 out of 10) throughout the experiment, which could indicate invalid responses and do not provide sufficient variance for analyses. Therefore, the final sample included 38 participants (Age: 18-51 (*M* = 28.5, *SD* = 8.19); Female = 17). This study was not preregistered.

#### Task

To examine the effects of challenge level on cognitive effort exertion and the associated affective experiences, we developed a self-paced incentivized cognitive control task based on one previously used by Leng and colleagues (Leng et al., 2021) (Figure 1A). In our task, participants were given fixed time intervals of 8 seconds to perform the classic color Stroop Task, which was designed to induce and measure cognitive effort (MacLeod, 1991). In the color-word Stroop task, participants had to name the ink color of a color word. There were four possible ink colors (red, yellow, green and blue) across four possible color words (‘RED’, ‘YELLOW’, ‘GREEN’, ‘BLUE’). Participants were instructed to press the key corresponding to the ink color of each stimulus. The ink color could be congruent (e.g., **BLUE**) or incongruent (e.g., **BLUE**) with the meaning of the word. Responding to incongruent stimuli has been shown to require an override of their more automatic tendency to respond based on the word meaning. The overall proportion of congruent (versus incongruent) trials was 50%. Due to the self-paced nature of design, the proportion of congruent trials could vary slightly across intervals, and was therefore included as a covariate in interval-level analyses. Participants were instructed to complete as many Stroop trials as they wanted during each time interval, and were told that each correct response would result in reward (in the form of “gems”), which could proportionately translate to monetary bonus rewards at the end of the task (5 gems = $0.01).

**Figure 1.**
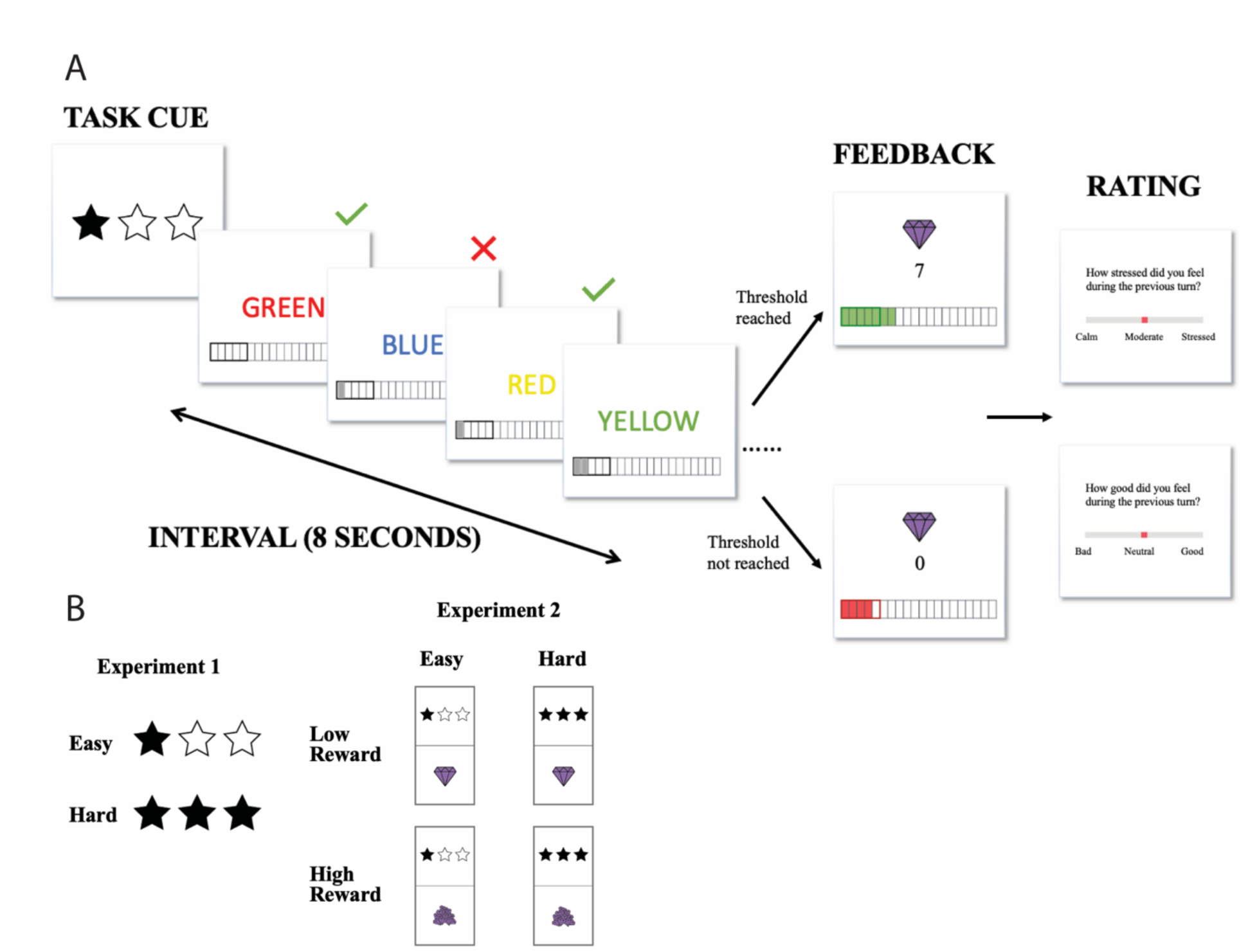
Task Schematics. (A) Each experiment contained 4 blocks of Stroop trials, with each block split into 8 smaller time intervals that lasted 8 seconds each. At the start of each interval, a cue is presented briefly to indicate challenge level (and reward level in Experiment 2) for the given interval. Participants can complete as many Stroop trials as they want before the time runs out. A tracker is displayed at the bottom of the screen and reflected the goal threshold and cumulative correct responses as real-time feedback in addition to the summary feedback at the end of each interval. If participants succeed in completing or exceeding the goal, they will have a chance to receive the reward corresponding to the number of trials they had completed. If participants failed to reach the goal, they will not receive any reward for that interval regardless of many correct responses they had made. After each interval, participants rated either their level of stress or positive affect during the preceding interval on a scale of 0 to 10. (B) Task cues in each experiment.

For each interval, there was a specific minimum number of correct responses that participants would have to complete in order to receive their rewards (the *goal threshold*). If the goal threshold was reached, the interval would yield a number of gems equal to the number of correct responses during that interval. If the goal threshold was not reached, the interval would yield 0 gems, regardless of the number of correct responses. We manipulated challenge level by varying the goal threshold across intervals. “Easy” intervals required only 5 correct responses to receive the bonus reward, whereas “Hard” required 8 correct responses to meet this threshold. These threshold values were selected based on pilot studies, which suggested that the higher goal threshold would generally be achievable but require more effort to meet than the lower goal threshold.

The challenge level for an upcoming interval was cued prior to the start of each interval (Figure 1B). A tracker at the bottom of the screen provided participants with real-time feedback regarding the cumulative number of correct trials within that interval. The tracker also served as a reminder of the goal threshold, indicating where their cumulative reward stood relative to the number of correct responses required. Participants also received feedback at the end of each interval regarding whether they had met their goal and how much reward (i.e., the number of gems) they earned for that interval.

We also measured affective experiences throughout the experiment. After each interval, we prompted participants to rate their affect during the preceding interval. Participants were asked to either rate their stress level (“How stressed did you feel during the previous turn?”) or their level of positive affect (“How good did you feel during the previous turn?”) on a scale of 0 to 10. Participants were only given one of these questions after each interval, with an equal number of each question asked and the ordering of the questions pseudorandomized within each block of 8 intervals.

The experiment was organized in large blocks of intervals of the same challenge level for within-subject comparisons of acute stress induced from said challenge. Challenge levels of blocks were varied pseudorandomly and in equal proportion across the experimental session. Each participant completed 4 blocks of 8 intervals per block and were instructed that the bonus reward from 2 intervals per block will be selected at the end of the task for bonus payment. This encouraged participants to treat each interval and block independently in terms of their level of effort investment, in part as a mitigant against fatigue effects, though all analyses also control for such order effects. The order of blocks was randomized across participants. Participants were also given ample practice with the Stroop task (score at least 5 correct trials in a row or a maximum of 60 trials) and task structure (4 intervals with the general interval structure and 2 intervals of practice with each set of cues) prior to the onset of the actual task. To ensure understanding of task instructions, participants were required to correctly complete short tests of task comprehension before continuing to the main experiment.

#### Measures and Analyses

As our main performance metrics, we measured reaction time (RT) and accuracy of Stroop trials. We also collected self-reported ratings of affect, as described above. With the current paradigm, we can analyze performance at the level of given time-intervals and at the level of individual trials of Stroop responses (Leng et al., 2021). We analyzed interval-level performance (correct trials per second) and self-report affective ratings by fitting linear mixed models (lme4 package in R; Bates et al., 2015) to estimate these parameters as functions of contrast-coded challenge level (Easy = -1, Hard = 1). The models controlled for the proportion of congruent vs. incongruent stimuli in a given interval, as well as the interval order within a block (1-8) and across the entire session (1-32). All continuous variables (e.g., interval order) were z-scored. All of our mixed models used maximally specified random effects (Barr et al., 2013).

We also analyzed accurate reaction time and accuracy at the trial level by fitting linear (RT) or logistic (accuracy) mixed models to estimate these parameters as functions of challenge level and whether the trial had been completed before or after the goal was reached. These models controlled for stimulus congruency, interval number, trial number over the course of the session, trial number within an interval, and a dummy variable of trial number within interval (- 1: first two trials within interval, 1: other trials). The last two trial number variables were added to account for variability in performance within an interval, which we describe in the Effects of goal completion and proximity on performance subsection of Results. Data and analysis code are available online (https://github.com/yzhangl/TSSS_Materials.git).

### Results

#### Effects of expected reward and challenge on overall performance

Overall, participants successfully met their minimum interval goals on 95.1% of easy intervals (goal = 5 correct trials) but only 74.8% of hard intervals (goal = 8 correct trials; *X^2^*(1, N = 38) = 13.54, *p* < 0.001, Table 1). On average, participants completed 8.62 correct trials per easy interval and 8.83 correct trials per hard interval. As an additional manipulation check, we examined whether participants had reached the high threshold goal (i.e., 8 cumulative correct trials) equally across conditions, especially since participants could have theoretically chosen to ignore the manipulation and instead respond as much as they can across both conditions. We found that participants reached the higher goal threshold on 69.5% of easy intervals, which is significantly lower than that in hard intervals (*X^2^*(1, 38) = 8.12, *p* = 0.004). This suggests that setting a challenge level affected how participants responded during the task.

**Table 1.** Mixed Model Results for Interval Goal Attainment and Higher Goal Attainment (Experiment 1)

| *Predictors* | Interval Goal Reached |  |  | High Goal Reached |  |  |
| --- | --- | --- | --- | --- | --- | --- |
|  | *Log Odds Ratios* | *SE* | *p-value* | *Log Odds Ratios* | *SE* | *p-value* |
| Hard - Easy | -1.15 | 0.10 | < **0.001\*\*\*** | 0.33 | 0.16 | **0.004\*\*\*** |
| Average Congruency | -0.21 | 0.10 | 0.080 | -0.15 | 0.08 | 0.089 |
| Scaled Interval Session Num<sup>1</sup> | 0.07 | 0.14 | 0.601 | -0.03 | 0.09 | 0.742 |
| Scaled Interval Block Num<sup>2</sup> | -0.06 | 0.11 | 0.627 | -0.12 | 0.08 | 0.155 |

1. Scaled interval number across the experiment session 2. Scaled interval number within each block

When participants faced hard intervals, they completed more correct trials per second (i.e., higher response rate) in a given interval compared with easy interval (*F*(1, 36.70) = 38.23, *p* < 0.001; Figure 2A, Table 2). This was reflected in faster trial-wise correct responses (i.e., speed, *F*(1, 35.8) = 22.73, *p* < 0.001; Figure 2B) and better trial accuracy (*X^2^*(1, N = 38) = 64.96, *p* < 0.001; Figure 2C) (Table 3). Notably, while response rate in easy intervals (*M* = 1.07, *SD* = 0.31) is lower on average compared with hard intervals (*M* = 1.10, *SD* = 0.30), it remains higher than expected, suggesting that participants continued to respond after the easy goal had been reached. In these models, we controlled for whether the participants had reached the interval goal, where we saw (as would be expected) higher response rate in intervals where the goal had been completed *F*(1, 36.7) = 421.89, *p* < 0.001), though the two constructs remain statistically dissociable (*r* = 0.62, *p* < 0.001). We again find this effect reflected in both faster speed of correct trial-wise responses (*F*(1, 26.5) = 79.18, *p* < 0.001) and better accuracy (*X*^2^(1, N = 38) = 226.71, *p* < 0.001) in intervals where the goal was reached versus not reached (Table 3). We note that while we controlled for within-interval variability in performance for reaction time and accuracy analyses, we were not able to do so for response rate given that the measure is a coarse estimation of performance at the interval level. However, all analyses were consistent and suggest that participants performed better in more challenging goals.

**Figure 2.**
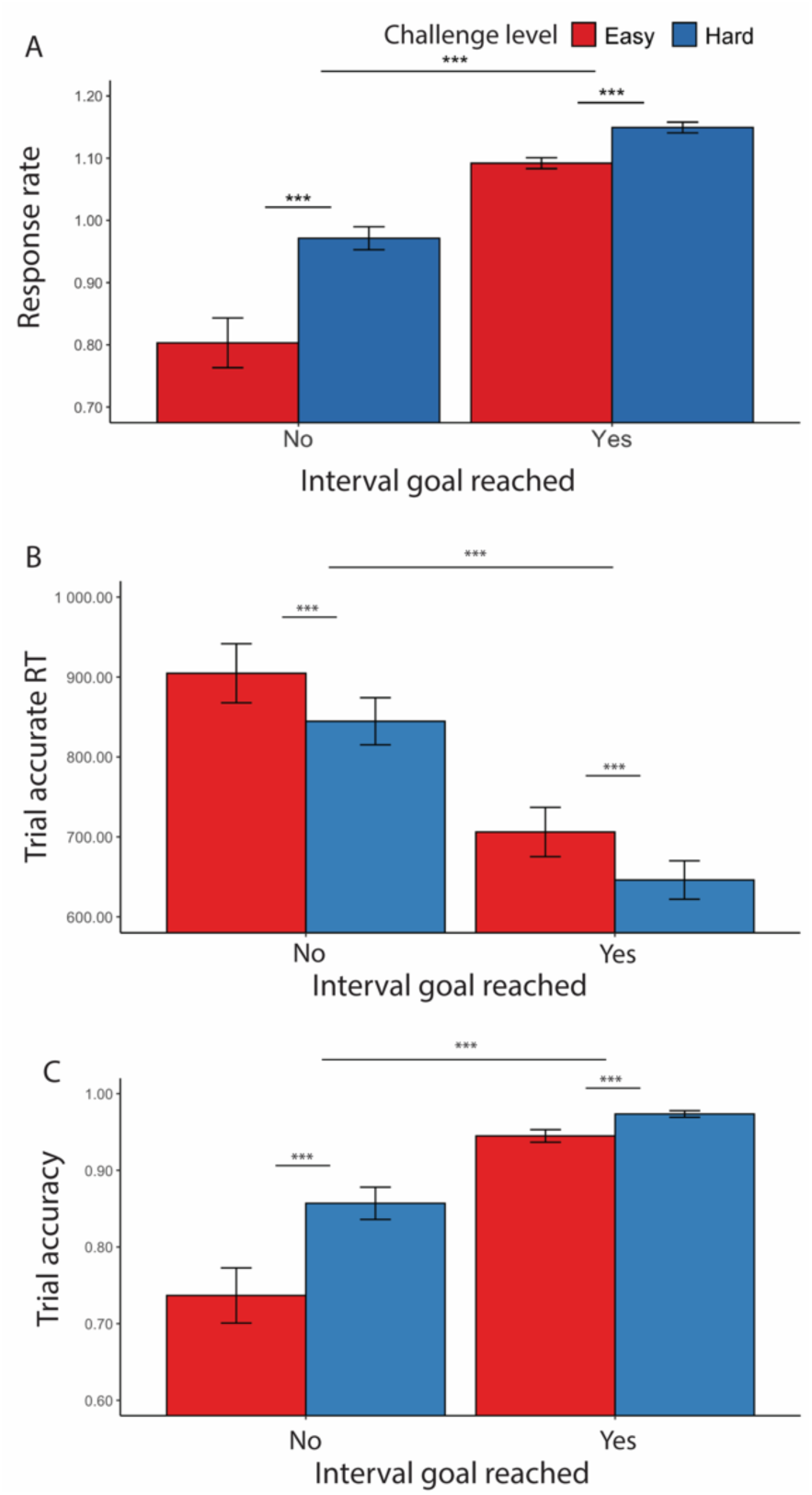
Effects of challenge level and goal completion on performance (Experiment 1). (A) When participants faced hard intervals, they completed more correct trials per second. When they had reached the interval goal, we found higher response rate compared with intervals where the goal was not reached. This is reflected in both (B) faster trial-wise reaction time for correct trials and (C) higher trial accuracy. Error bars reflect standard errors. Asterisks denote the significance level of main effects. ***: *p* < 0.001

**Table 2.** Mixed Model Results for Correct Responses per Second (Experiment 1)

| *Predictors* | Response Rate |  |  |
| --- | --- | --- | --- |
|  | *Estimates* | *SE* | *p-value* |
| Hard - Easy | 0.05 | 0.01 | < **0.001\*\*\*** |
| Interval Goal Completion<sup>1</sup> | 0.42 | 0.02 | < **0.001\*\*\*** |
| Average Congruency | 0.00 | 0.01 | 0.614 |
| Scaled Interval Session Num | -0.00 | 0.01 | 0.336 |
| Scaled Interval Block Num | 0.01 | 0.00 | 0.508 |

1. Whether participants had reached the goal for that interval (1: reached, 0: not reached)

**Table 3.**
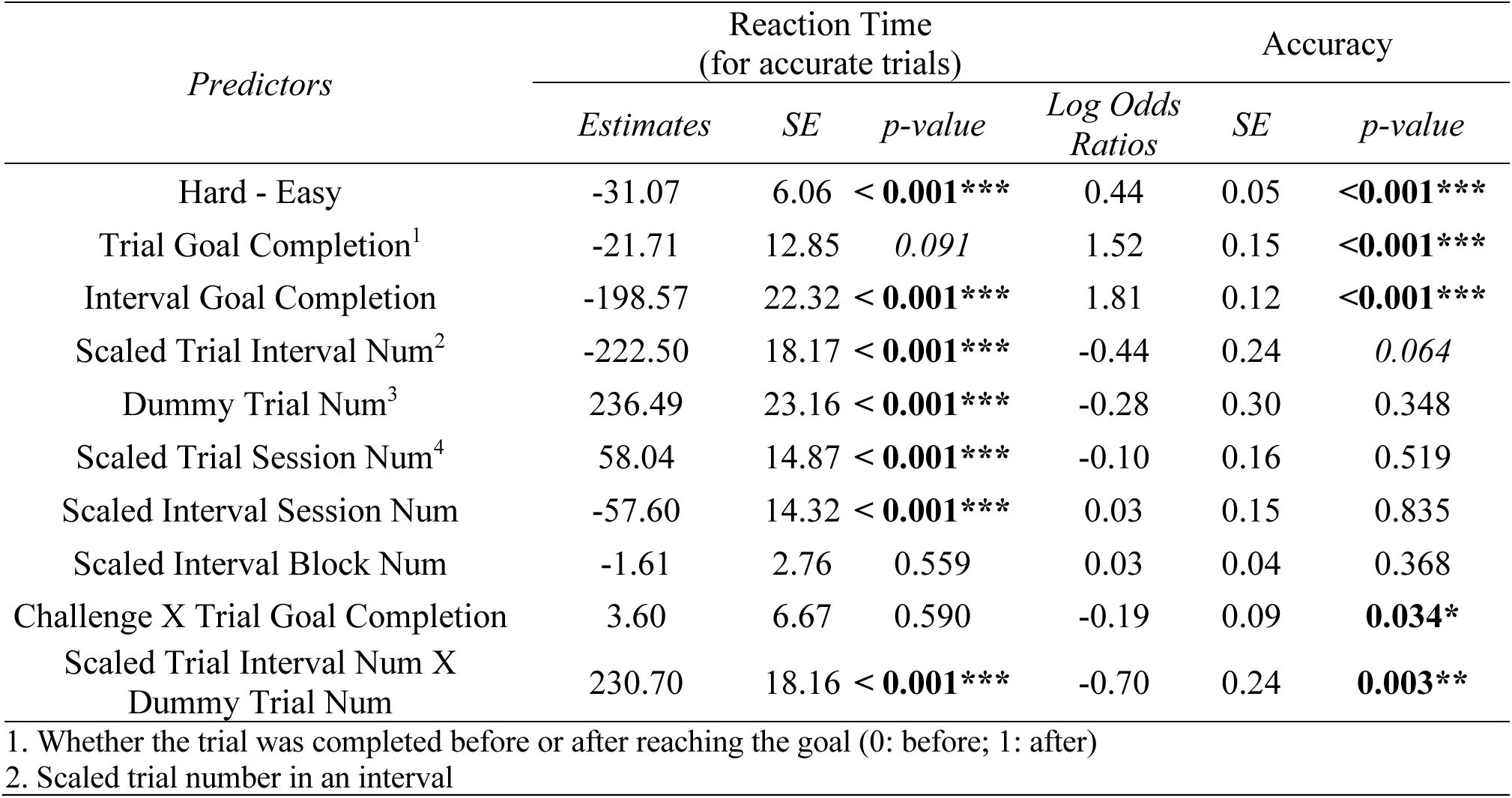
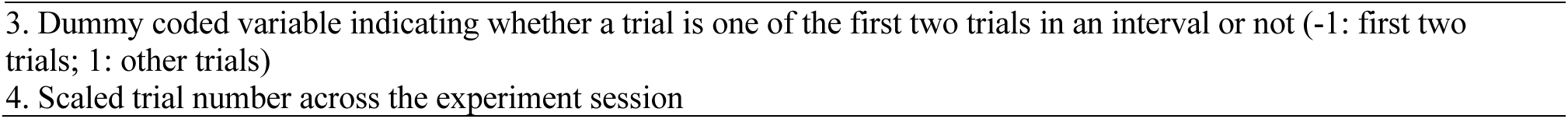
Mixed Model Results for Reaction Time and Accuracy (Experiment 1)

| *Predictors* | Reaction Time<br>(for accurate trials) |  |  | Accuracy |  |  |
| --- | --- | --- | --- | --- | --- | --- |
|  | *Estimates* | *SE* | *p-value* | *Log Odds Ratios* | *SE* | *p-value* |
| Hard - Easy | -31.07 | 6.06 | < **0.001\*\*\*** | 0.44 | 0.05 | <**0.001\*\*\*** |
| Trial Goal Completion<sup>1</sup> | -21.71 | 12.85 | 0.091 | 1.52 | 0.15 | <**0.001\*\*\*** |
| Interval Goal Completion | -198.57 | 22.32 | < **0.001\*\*\*** | 1.81 | 0.12 | <**0.001\*\*\*** |
| Scaled Trial Interval Num<sup>2</sup> | -222.50 | 18.17 | < **0.001\*\*\*** | -0.44 | 0.24 | 0.064 |
| Dummy Trial Num<sup>3</sup> | 236.49 | 23.16 | < **0.001\*\*\*** | -0.28 | 0.30 | 0.348 |
| Scaled Trial Session Num<sup>4</sup> | 58.04 | 14.87 | < **0.001\*\*\*** | -0.10 | 0.16 | 0.519 |
| Scaled Interval Session Num | -57.60 | 14.32 | < **0.001\*\*\*** | 0.03 | 0.15 | 0.835 |
| Scaled Interval Block Num | -1.61 | 2.76 | 0.559 | 0.03 | 0.04 | 0.368 |
| Challenge X Trial Goal Completion | 3.60 | 6.67 | 0.590 | -0.19 | 0.09 | **0.034\*** |
| Scaled Trial Interval Num X<br>Dummy Trial Num | 230.70 | 18.16 | < **0.001\*\*\*** | -0.70 | 0.24 | **0.003\*\*** |

1. Whether the trial was completed before or after reaching the goal (0: before; 1: after) 2. Scaled trial number in an interval
- 3. Dummy coded variable indicating whether a trial is one of the first two trials in an interval or not (-1: first two trials; 1: other trials) 4. Scaled trial number across the experiment session -

#### Effects of goal completion and proximity on performance

The findings above describe aggregate performance as challenge level varied, but our experimental design allowed us to examine performance on a more granular level: to see how it varied on a trial-by-trial level as participants approached the goal, and after they surpassed it. In our task, prior to reaching the minimum number of trials for a given interval, participants were incentivized to reach the goal before the deadline (otherwise risking foregoing any reward for that interval). After meeting this minimum goal, though, they were still rewarded for each correct response and were therefore incentivized to keep completing as many trials as they could, such that participants could obtain the same amount of reward across both easy and hard intervals. We were therefore interested in the extent to which performance would maintain or differ before and after reaching a goal (focusing only on intervals in which that minimum goal was met). These analyses additionally controlled for variability in performance within an interval.

Consistent with the findings above, we found a main effect of challenge level on performance, such that participants were more likely to respond accurately (*F*(1, 25.7) = 27.42, *p* < 0.001), and faster to do so (*X^2^*(1, N = 38) = 34.98, *p* < 0.001), during more challenging intervals (Figure 3, Table 4). We further found that performance differed before versus after meeting the goal, with trials completed after meeting an interval’s threshold being faster (*F*(1, 59.4) = 6.97, *p* = 0.008; Figure 3A) and more accurate (*X^2^*(1, N = 38) = 84.96, *p* < 0.001; Figure 3B) (Table 4).

**Figure 3.**
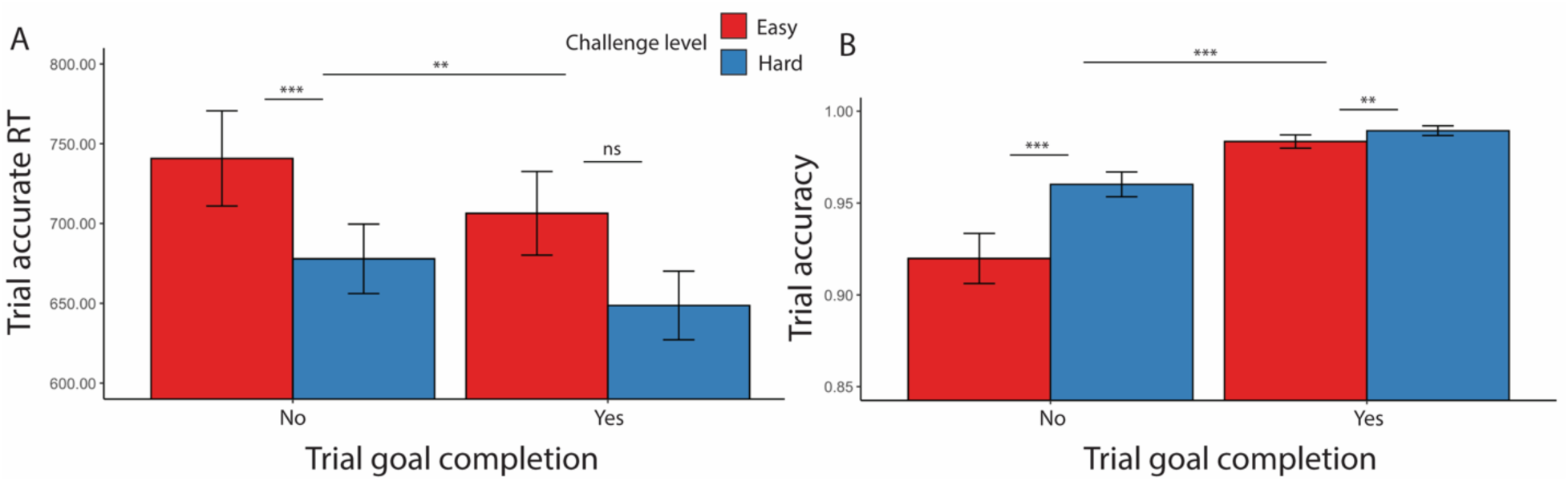
Goal completion effects (goal reached intervals only, Experiment 1). (A) Participants were overall faster in completing accurate trials and (B) more accurate after they had reached the interval goal. Error bars reflect standard errors. Asterisks denote the significance level of main effects. ***: *p* < 0.001; **: *p* < 0.01

**Table 4.** Mixed Model Results for Reaction Time and Accuracy (Experiment 1, goal-reached intervals only)

| *Predictors* | Reaction Time<br>(for accurate trials) |  |  | Accuracy |  |  |
| --- | --- | --- | --- | --- | --- | --- |
|  | *Estimates* | *SE* | *p-value* | *Log Odds Ratios* | *SE* | *p-value* |
| Hard - Easy | -31.45 | 6.01 | < **0.001** *** | 0.37 | 0.06 | < **0.001** *** |
| Trial Goal Completion<sup>1</sup> | -31.81 | 12.05 | **0.008** ** | 1.50 | 0.16 | < **0.001** *** |
| Scaled Trial Interval Num | -166.37 | 16.30 | < **0.001** *** | -0.26 | 0.28 | 0.355 |
| Dummy Trial Num | 177.71 | 20.78 | < **0.001** *** | -0.43 | 0.36 | 0.232 |
| Scaled Trial Session Num | 7.42 | 14.29 | 0.603 | -0.28 | 0.20 | 0.171 |
| Scaled Interval Session Num | -4.25 | 14.07 | 0.763 | 0.22 | 0.20 | 0.291 |
| Scaled Interval Block Num | -1.25 | 2.41 | 0.604 | 0.04 | 0.04 | 0.369 |
| Challenge X Trial Goal Completion | 2.57 | 5.58 | 0.645 | -0.15 | 0.09 | 0.113 |
| Scaled Trial Interval Num X Dummy Trial Num | 181.08 | 16.30 | < **0.001** *** | -0.89 | 0.28 | **0.001** *** |

1. Whether the trial was completed before or after reaching the goal (0: before; 1: after) 2. Scaled trial number across the experiment session

We also examined whether goal proximity, namely how far a certain trial was from the goal threshold, had an impact on performance. We similarly selected for intervals where the minimum goal had been completed, and further controlled for an interval’s challenge level. We found that participants initially sped up after starting the interval for approximately 2 trials, then gradually slowed as they neared and surpassed the goal (Figure 4). To avoid conflating the effect of goal proximity with distance from this initial speeding effect, we excluded those first two trials in each interval. We also excluded the last trial in each interval to avoid potential confounds related to unstable performance at the end of each time interval. After excluding these trials, we found that participants slowed down in making correct responses as they neared the goal (*F*(1, 23.5) = 34.32, *p* < 0.001), while maintaining similar levels of accuracy (*X^2^*(1, N = 38) = 0.07, *p* = 0.850, Table 5, Figure 4). We also found an interaction between goal proximity and challenge level on accuracy, such that participants became less accurate when nearing a challenging goal, but not when nearing the easier goal (*X^2^*(1, N = 38) = 7.20, *p* = 0.007). While our analyses control for order effects across the session (e.g., interval and block number), we cannot rule out the possibility that this interaction reflects the fact that goal-proximal trials on challenging intervals are also later in the interval, and potentially reflect within-interval order effects such as fatigue (note that the same concern does not hold for the main effects reported above).

**Figure 4.**
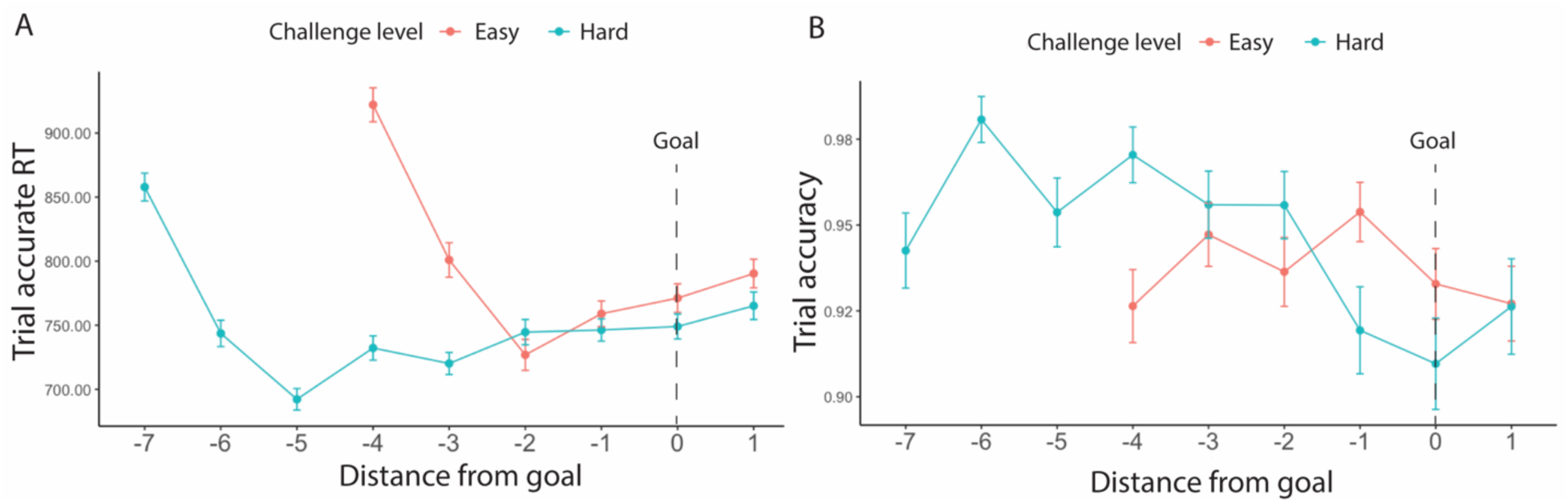
Goal proximity effects (Experiment 1). Overall, participants were slowest early in the interval (during the first two trials) and then gradually slowed again as they got closer to reaching their goal. Their accuracy did not vary with goal proximity. Error bars reflect standard errors.

**Table 5.** Mixed Model Results for Goal Proximity Effect for Reaction Time and Accuracy (Experiment 1)

| Predictors | Response Time<br>(for accurate trials) |  |  | Accuracy |  |  |
| --- | --- | --- | --- | --- | --- | --- |
|  | Estimates | SE | p-value | Log Odds Ratios | SE | p-value |
| Goal Distance<sup>1</sup> | -27.88 | 4.76 | < **0.001\*\*\*** | -0.01 | 0.07 | 0.850 |
| Hard – Easy | -14.87 | 4.62 | **0.001\*\*\*** | 0.12 | 0.08 | 0.102 |
| Scaled Trial Session Num | 3.65 | 15.47 | 0.813 | -0.49 | 0.13 | **0.019\*** |
| Scaled Interval Session Num | -2.47 | 15.42 | 0.873 | 0.37 | 0.31 | 0.089 |
| Scaled Interval Block Num | -4.40 | 2.62 | 0.093 | 0.08 | 0.06 | 0.118 |
| Goal Distance X Challenge | 3.47 | 3.28 | 0.291 | 0.17 | 0.08 | **0.007\*\*** |

Together, these results show that more challenging goals motivate better overall performance, seen in both faster and more accurate responses. Within a given interval, we see that these challenge-related performance improvements are reflected in trials both before and after goal completion.

#### Effects of challenge level and performance on affective experiences

We next examined what elements of the task led to changes in affective states. As expected, we found that one’s success or failure at reaching their goal for a given threshold significantly influenced affect (Figure 5, Table 6): participants reported feeling less positive (*F*(1, 19.6) = 26.73, *p* < 0.001) and more stressed (*F*(1, 54.1) = 29.18, *p* < 0.001) during intervals where they failed to reach a goal.

**Figure 5.**
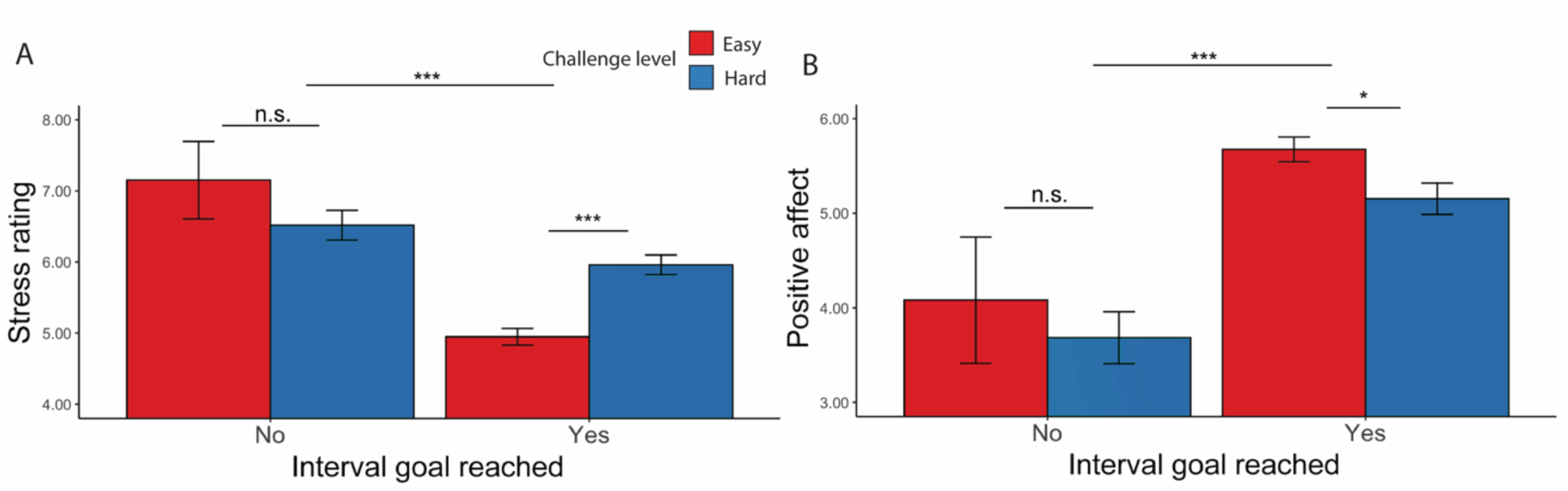
Affective Ratings based on goal completion and challenge level (Experiment 1). (A) Stress Ratings: Participants reported feeling more stress when they failed to reach the goal. In intervals where the goal was reached, they also reported feeling more stressed in hard intervals. (B) Positive Affect: Along the same line, participants reported to have felt worse when they failed to complete the goal and after completing the goal in hard intervals. Error bars reflect standard errors. Asterisks denote the significance level of main effects. ***: *p* < 0.001; *: *p* < 0.05

**Table 6.**
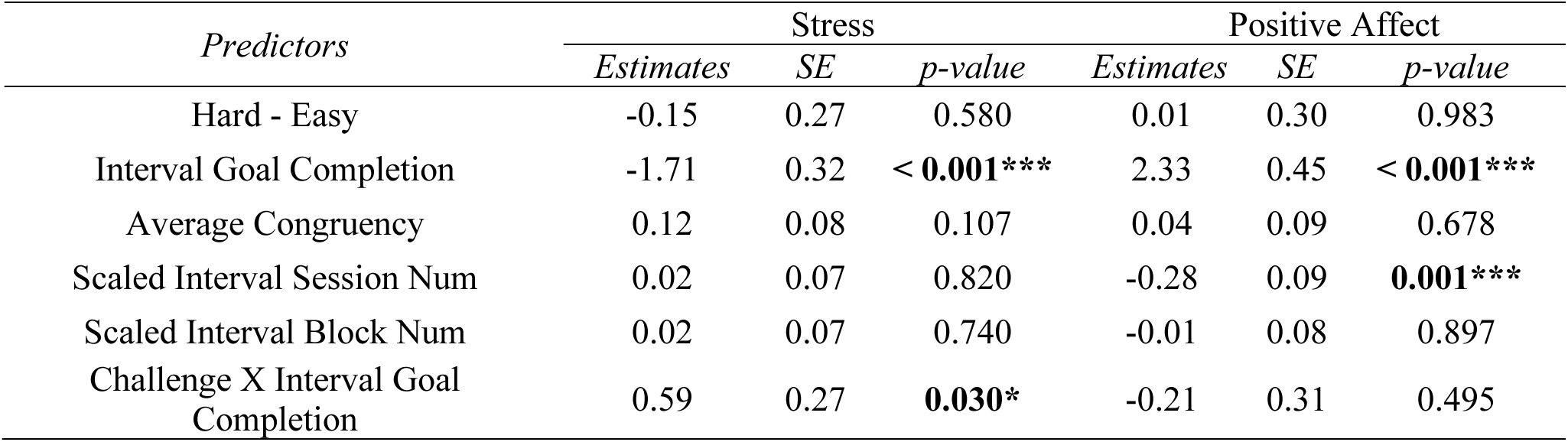
Mixed Model Results for Affective Ratings Based on Challenge Level and Goal Completion (Experiment 1)

| *Predictors* | Stress |  |  | Positive Affect |  |  |
| --- | --- | --- | --- | --- | --- | --- |
|  | *Estimates* | *SE* | *p-value* | *Estimates* | *SE* | *p-value* |
| Hard - Easy | -0.15 | 0.27 | 0.580 | 0.01 | 0.30 | 0.983 |
| Interval Goal Completion | -1.71 | 0.32 | < **0.001\*\*\*** | 2.33 | 0.45 | < **0.001\*\*\*** |
| Average Congruency | 0.12 | 0.08 | 0.107 | 0.04 | 0.09 | 0.678 |
| Scaled Interval Session Num | 0.02 | 0.07 | 0.820 | -0.28 | 0.09 | **0.001\*\*\*** |
| Scaled Interval Block Num | 0.02 | 0.07 | 0.740 | -0.01 | 0.08 | 0.897 |
| Challenge X Interval Goal Completion | 0.59 | 0.27 | **0.030\*** | -0.21 | 0.31 | 0.495 |

Focusing on intervals where participants had successfully met their goal, we found main effects of challenge level on affective experiences, such that participants felt worse (*F*(1, 38.5) = 5.19, *p* = 0.023) and more stressed (*F*(1, 35.3) = 14.44, *p* < 0.001) while performing hard intervals compared with easy intervals (Table 7). Similarly in intervals where the goal was completed and reward was given, we found that when participants completed more correct responses per second, they reported less stress (*F*(1, 39.0) = 24.47, *p* < 0.001) and greater positive affect (*F*(1, 38.8) = 36.72, *p* < 0.001; Table 8, Figure 6A-B). This was reflected in analogous associations with faster average reaction time of correct trials (less stress: *F*(1, 39.83) = 7.75, *p* = 0.006; greater positive affect: *F*(1, 25.14) = 10.30, *p* = 0.001; Table 9, Figure 6C-D) and higher average accuracy (less stress: *F*(1, 34.33) = 7.75, *p* < 0.001; greater positive affect: *F*(1, 37.39) = 41.54, *p* < 0.001; Table 10, Figure 6E-F).

**Figure 6.**
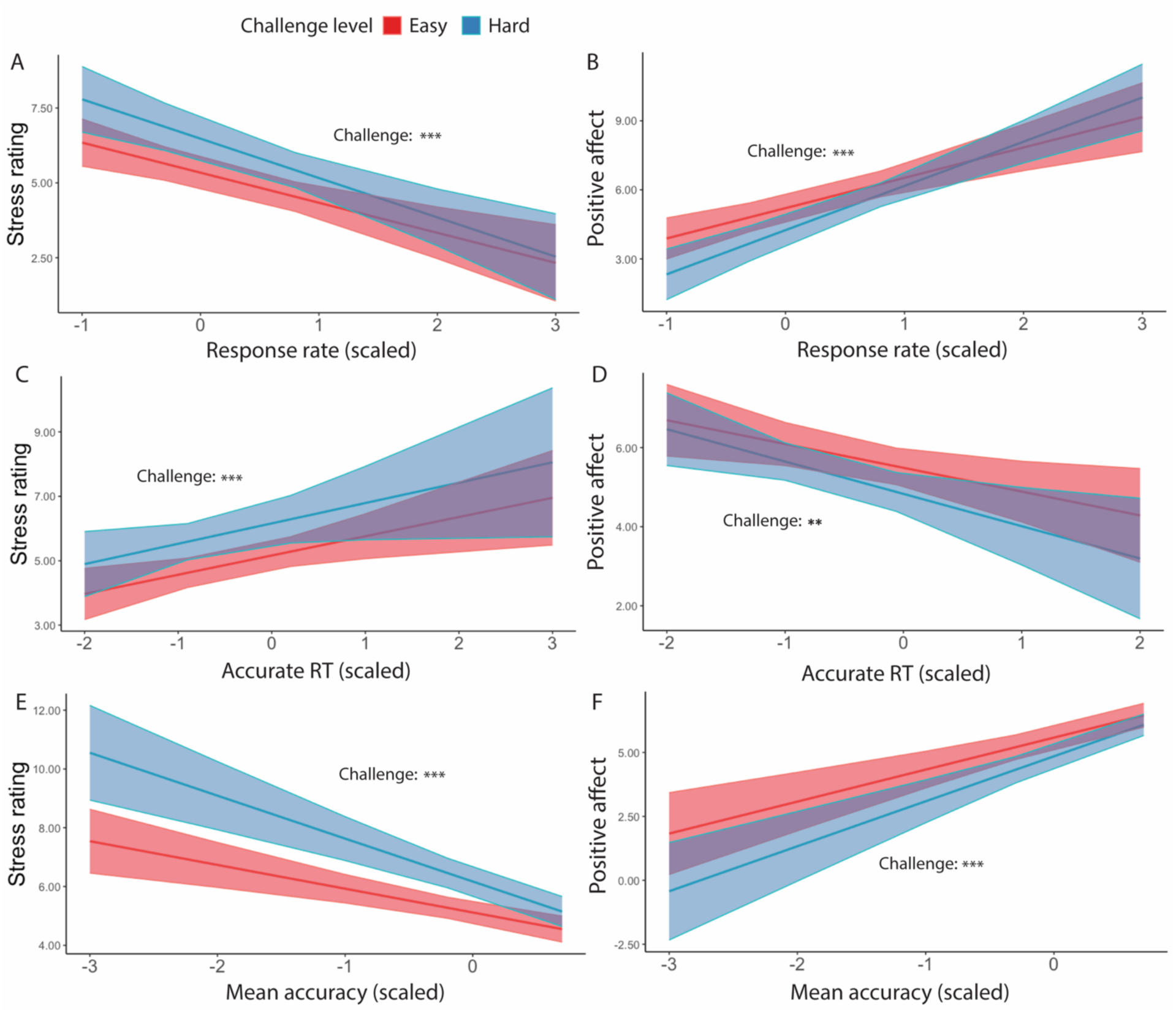
Affective Ratings predicted by performance, for goal-completed intervals only (Experiment 1). We found that in intervals where the goal had been reached, participants reported to have felt less stressed when (A) performed more correct trials per interval, (C) faster reaction time, and (E) higher accuracy. They also reported feeling better overall when (B) performed more correct trials per interval, (D) faster reaction time, and (F) higher accuracy. Error bars reflect standard errors. Asterisks denote the significance level of main effects. ***: *p* < 0.001; **: *p* < 0.01

**Table 7.** Mixed Model Results for Affective Ratings Based on Challenge Level (Goal Completed Intervals Only, Experiment 1)

| *Predictors* | Stress |  |  | Positive Affect |  |  |
| --- | --- | --- | --- | --- | --- | --- |
|  | *Estimates* | *SE* | *p-value* | *Estimates* | *SE* | *p-value* |
| Hard - Easy | 0.46 | 0.12 | < **0.001** *** | -0.23 | 0.10 | **0.023** * |
| Average Congruency | 0.12 | 0.09 | 0.159 | 0.01 | 0.10 | 0.947 |
| Scaled Interval Session Num | -0.02 | 0.08 | 0.782 | -0.35 | 0.09 | < **0.001** *** |
| Scaled Interval Block Num | 0.03 | 0.08 | 0.746 | 0.08 | 0.09 | 0.356 |

**Table 8.** Mixed Model Results for Affective Ratings Based on Response Rate (Goal Completed Intervals Only, Experiment 1)

| *Predictors* | Stress |  |  | Positive Affect |  |  |
| --- | --- | --- | --- | --- | --- | --- |
|  | *Estimates* | *SE* | *p-value* | *Estimates* | *SE* | *p-value* |
| Response Rate | -1.16 | 0.23 | < **0.001** *** | 1.62 | 0.27 | < **0.001** *** |
| Hard - Easy | 0.57 | 0.11 | < **0.001** *** | -0.48 | 0.09 | < **0.001** *** |
| Average Congruency | 0.13 | 0.07 | 0.084 | -0.06 | 0.07 | 0.390 |
| Scaled Interval Session Num | -0.02 | 0.07 | 0.780 | -0.32 | 0.07 | < **0.001** *** |
| Scaled Interval Block Num | 0.05 | 0.07 | 0.500 | -0.03 | 0.07 | 0.697 |
| Response Rate X Challenge | -0.15 | 0.11 | 0.172 | 0.30 | 0.10 | **0.002** ** |

**Table 9.** Mixed Model Results for Affective Ratings Based on Average Reaction Time (Goal Completed Intervals Only, Experiment 1)

| *Predictors* | Stress |  |  | Positive Affect |  |  |
| --- | --- | --- | --- | --- | --- | --- |
|  | *Estimates* | *SE* | *p-value* | *Estimates* | *SE* | *p-value* |
| Mean Accurate RT | 0.61 | 0.22 | **0.006\*\*** | -0.71 | 0.22 | **0.001\*\*\*** |
| Hard - Easy | 0.50 | 0.14 | **< 0.001\*\*\*** | -0.33 | 0.12 | **0.004\*\*** |
| Average Congruency | 0.15 | 0.08 | 0.074 | -0.03 | 0.10 | 0.721 |
| Scaled Interval Session Num | -0.04 | 0.08 | 0.646 | -0.35 | 0.09 | **< 0.001\*\*\*** |
| Scaled Interval Block Num | 0.01 | 0.08 | 0.854 | 0.06 | 0.09 | 0.472 |
| Mean Accurate RT X Challenge | 0.02 | 0.16 | 0.911 | -0.11 | 0.15 | 0.473 |

**Table 10.** Mixed Model Results for Affective Ratings Based on Average Accuracy (Goal Completed Intervals Only, Experiment 1)

| *Predictors* | Stress |  |  | Positive Affect |  |  |
| --- | --- | --- | --- | --- | --- | --- |
|  | *Estimates* | *SE* | *p-value* | *Estimates* | *SE* | *p-value* |
| Mean Accuracy | -1.13 | 0.17 | **< 0.001\*\*\*** | 1.51 | 0.23 | **< 0.001\*\*\*** |
| Hard - Easy | 0.53 | 0.10 | **< 0.001\*\*\*** | -0.36 | 0.09 | **< 0.001\*\*\*** |
| Average Congruency | 0.07 | 0.08 | 0.387 | 0.00 | 0.08 | 0.988 |
| Scaled Interval Session Num | -0.02 | 0.07 | 0.740 | -0.32 | 0.08 | **< 0.001\*\*\*** |
| Scaled Interval Block Num | 0.04 | 0.08 | 0.561 | 0.02 | 0.07 | 0.76 |
| Mean Accuracy X Challenge | -0.33 | 0.13 | **0.010\*\*** | 0.25 | 0.12 | **0.040\*** |

We also observed a significant interaction between challenge level and response rate in predicting positive affect (*F*(1, 149.1) = 9.69, *p* = 0.002), such that the increase in positive affect due to higher response rate was enhanced in more challenging relative to less challenging intervals. Similar interactions were observed between average accuracy and challenge level in predicting stress ratings (*F*(1, 395.3) = 6.62, *p* = 0.010) and positive affect (*F*(1, 38) = 4.21, *p* = 0.040), such that performing with higher accuracy reduced the challenge-induced affective experience (i.e., higher stress, less positive affect). This interaction between performance and challenge level was only observed for accuracy but not correct response time (*p*s > 0.470). These results suggest that the challenge-induced affective experience (i.e., higher stress, less positive affect) was reduced as participants performed better.

Taken together, these results indicate that challenge level and performance both impact affective experiences during the task, such that easier intervals and better performance were associated with more positive and less negative affect. Notably, better performance, particularly higher accuracy, appeared to mitigate the negative affect that was brought about by challenge level (provided the participant had met the goal for that interval).

### Experiment 1 Discussion

Experiment 1 provided evidence that our manipulation of expected challenge level improved performance (i.e., faster and more accurate responses), such that attempting to reach a harder goal motivated people to exert more cognitive effort. Though participants could have chosen to ignore the thresholds and attempt equally hard across challenge levels to gain as much reward bonus as possible, our results are consistent with behavioral patterns that reflect an increase in selective attention to task in past research and could potentially suggest that more challenging condition leads participants to allocate more attention on the task at hand (Leng et al., 2021). Our findings over the course of a given interval are further in line with this, with faster and more accurate performance after goal completion, indicating that participants not only tried harder to achieve their goal in hard intervals, but also persisted with higher levels of effort after reaching that minimum goal. This increase in performance could reflect immediate relief felt by participants when the stakes of completing more correct trials decreased significantly after reaching the goal. Results could also be interpreted in terms of parameters of sequential sampling models, which suggests that noisy evidence favoring each alternative response is integrated over time and a response is made when sufficient evidence has accumulated favoring one alternative over the other (Bogacz et al., 2006). The rate of evidence is drift rate while the amount of evidence needed for the initiation of response is boundary separation. The behavioral patterns we observed have also been found to suggest greater selective attention to task (i.e., greater evidence accumulation rate) and lower response caution (i.e., lower response threshold) (Leng et al., 2021). Taken together, these results are consistent with current work, such that a specific and challenging goal motivates the output of more cognitive effort and prolonged effort exertion compared with an easy goal.

Due to the structure of our task, we were able to examine performance with respect to the distance from the goal. We found that participants slowed down on trials near the goal while performing with similar accuracy, which could potentially indicate reduced rate of evidence accumulation (i.e., reduced selective attention) and higher response threshold (i.e., increased response caution) as they approached the goal (Leng et al., 2021). We also found that greater proximity to the goal interacted with challenge level to further boost accuracy on the task, potentially suggesting that closer distance to the goal provides additional motivation to complete more challenging goals relative to less challenging ones. It is also possible that the interaction with challenge level is because goal completion for the harder challenge level requires longer effort exertion and occurs later in the trial (i.e., greater time constraints), making it more difficult to relax immediately after reaching the goal compared with easy intervals. We also cannot rule out the possibility that participants’ fatigue within each interval contributed to lower accuracy differences between challenge levels so these results should be interpreted with caution. With our current experimental design, we are unable to address this limitation of time constraints in data analyses with certainty. Future studies could do so by collecting the time point at which participants reach the goal.

Our results also support the hypothesis that challenge level and performance both play a role in influencing affective experiences during the task. We found that ratings of both stress and positive affect were predominantly driven by goal completion, with less stress and higher positive affect when the goal had been completed. When focusing on the majority of intervals where participants were successful in meeting the goal, affective ratings were primarily influenced by challenge level. More challenging goals led participants to feel more stressed and less positive while performing the task, which was not mitigated by eventually reaching the goal. Interestingly, we found that performance and challenge level interacted in predicting affect, such that higher accuracy had a larger impact on affective states when the goal was more challenging (for instance, leading them to feel greater relief at reaching one’s goal), consistent with the broader hypothesis that the effects of performance and challenge level on affective states are intertwined.

While these findings address the role of challenge level in shaping performance and affect, they leave open the question of how these variables are additionally shaped by the rewards at stake (which were held constant in this experiment). Whereas previous work suggests that higher rewards should motivate better performance on this task (Leng et al., 2021), whether these reward effects will be exacerbated or diminished by challenge level is unclear, as is the question of how reward and challenge level will separately and interactively influence affective experiences in this task. Experiment 2 sought to examine these questions by varying both the challenge level and the size of the incentives for correct performance.

## Experiment 2: The integrative influence of challenge level and reward incentives on performance and affect

### Methods

#### Participants

We recruited 79 participants online through Prolific using the same criteria as Experiment Consent and IRB approval was given, and monetary reward and bonus were received for participation. One participant was excluded in the interval-level analysis due to a lack of variance in their affective ratings throughout the experiment, leaving a total of 78 participants (45 Female) for analyses, aged 18-53 (*M* = 27.21, *SD* = 8.89)

#### Task

In addition to varying challenge levels, we also varied reward levels in this experiment. Similar to Experiment 1, we used an interval-based task structure to measure cognitive effort persistence. Goal thresholds were set up in the same manner, such that a number of cumulative correct responses were needed in an interval to reach the goal (5 for easy, 8 for hard). In addition, participants were instructed that intervals varied in their reward values. In Low Reward intervals, each correct response earned 1 gem, whereas High Reward correct responses gave 10 gems. Therefore, the task included four conditions (Low Easy, Low Hard, High Easy, High Hard).

The task was grouped by blocks, where there were 8 blocks of 8 intervals each. Each block contained intervals with only one type of challenge level and semi-randomized reward types. Participants were again instructed that two intervals from each block would be chosen for bonus payment. Stress and positive affect were measured with the same self-reports of affective ratings as in Experiment 1. Similar to Experiment 1, we analyzed interval-level performance (correct trials per second) and self-report affective ratings by fitting linear mixed models (lme4 package in R; Bates et al., 2015) to estimate these parameters as functions of contrast-coded challenge level (Easy = -1, Hard = 1) and reward level (Low Reward = -1, High Reward = 1), and their interactions.

### Results

#### Effects of expected reward and challenge on overall performance

Overall, participants met the minimum interval goals on 96.6% of easy intervals (average of 8.40 trials per interval) and 73.4% of hard intervals (8.67 trials) (*X^2^*(1, N = 78) = 92.47, *p* < 0.001; Table 11), which were very similar to Experiment 1. As in Experiment 1, we also found that participants reached the higher goal threshold less often (66.9% of intervals) for easy intervals than for hard intervals (73.4%; *X^2^*(1, N = 78) = 23.78, *p* < 0.001). Building on these results, we found that participants were just as likely to reach the minimum goal for a given interval when the reward for each correct response was low (84.8%; average of 8.46 trials) as when it was high (85.1%; 8.61 trials) (*p* = 0.983). However, higher rewards motivated participants to reach the *higher* threshold more often (71.6%) than low rewards (68.7%) (*X^2^*(1, N = 78) = 6.05, *p* = 0.019).

**Table 11.** Mixed Model Results for Interval Goal Attainment and Higher Goal Attainment Based on Challenge and Reward Level (Experiment 2)

| *Predictors* | Interval Goal Reached |  |  | High Goal Reached |  |  |
| --- | --- | --- | --- | --- | --- | --- |
|  | *Log Odds Ratios* | *SE* | *p-value* | *Log Odds Ratios* | *SE* | *p-value* |
| Hard - Easy | -1.52 | 0.16 | < **0.001\*\*\*** | 0.22 | 0.05 | < **0.001\*\*\*** |
| High Reward – Low Reward | 0.00 | 0.09 | 0.983 | 0.10 | 0.04 | **0.019\*** |
| Average Congruency | 0.07 | 0.05 | 0.150 | 0.10 | 0.04 | **0.011\*** |
| Scaled Interval Session Num | 0.17 | 0.05 | **0.001\*\*\*** | 0.17 | 0.04 | < **0.001\*\*\*** |
| Scaled Interval Block Num | 0.02 | 0.05 | 0.676 | 0.05 | 0.04 | 0.167 |
| Challenge X Reward | 0.06 | 0.07 | 0.396 | -0.05 | 0.04 | 0.179 |

Consistent with Experiment 1, we found that participants completed more correct trials per second when faced with hard intervals compared with easy intervals (*F*(1, 77.1) = 142.53, *p* < 0.001; Table 12, Figure 7A), which was reflected in faster trial-wise correct responses (*F*(1, 77) = 62.27, *p* < 0.001, Figure 7B) and better trial-level accuracy (*X^2^*(1, N=78) = 215.12, *p* < 0.001, Figure 7C) (Table 13). Also similar to Experiment 1, response rate for easy intervals (*M* = 1.05, *SD* = 0.26) remained high but is lower compared with that in hard intervals (*M* = 1.09, *SD* = 0.26). Participants also exhibited higher response rates when faced with larger potential rewards (*F*(1, 3975.8) = 16.83, *p* < 0.001, Table 12). These effects of reward and challenge level on performance appear to be independent and additive, as we did not observe an interaction (*F*(1, 4809.6) = 0.94, *p* = 0.333). These results remained when controlling for whether the interval goal had been reached (Table 13, Figure 7B-C). Similar to Experiment 1, response rate was higher when the interval goal was completed but the two constructs remained dissociable (*r* = 0.56, *p* < 0.001). When controlling for these variables, we found that participants were faster to complete correct trials in high relative to low reward intervals (*F*(1, 1252) = 10.18, *p* = 0.001) without differing in their overall accuracy (*X^2^*(1, N =78) = 0.16, *p* = 0.693) (Table 13).

**Figure 7.**
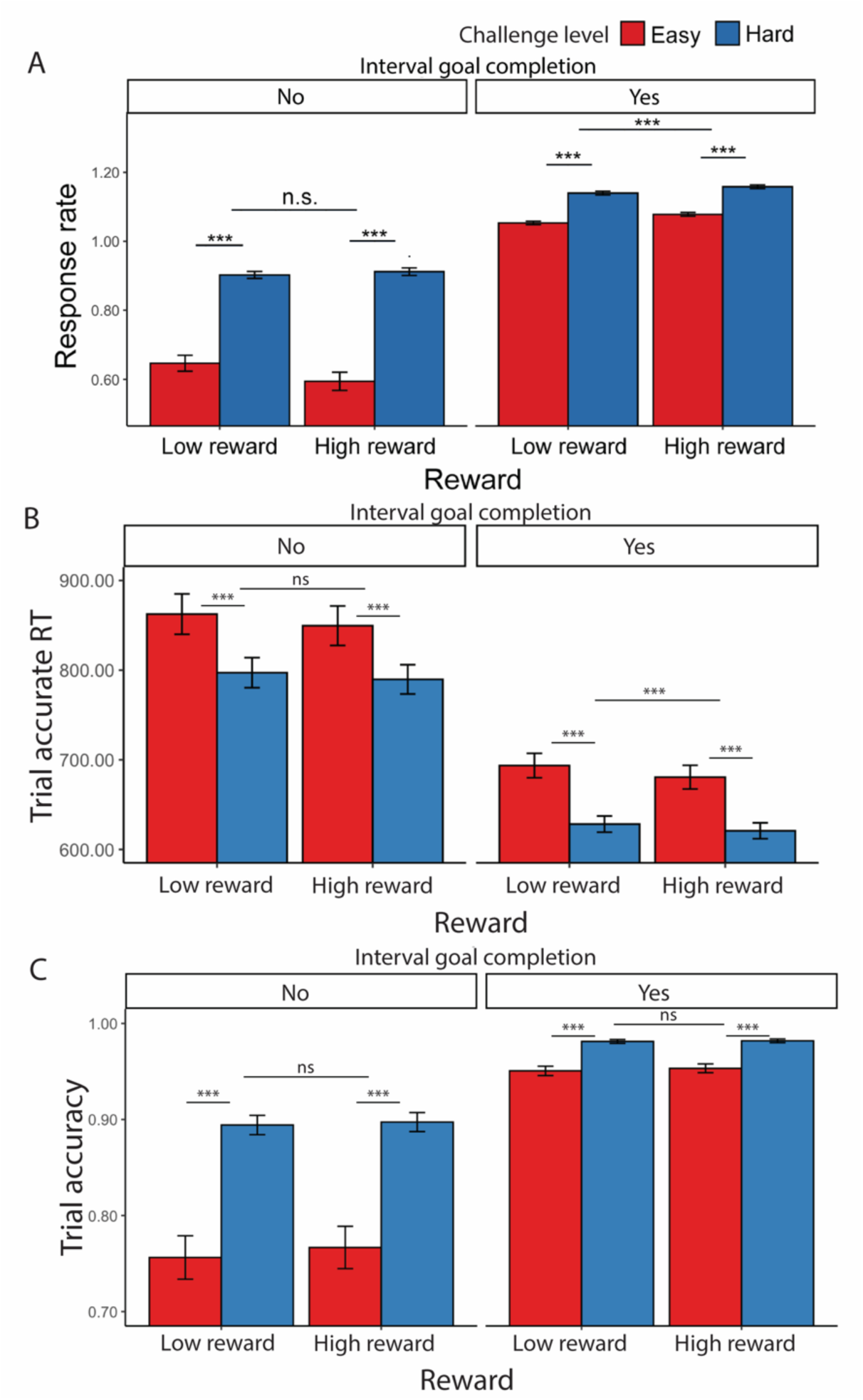
Effects of challenge level, reward level, and goal completion on performance. (A) When participants faced hard intervals and when there are higher incentives, they completed more correct trials per second. When they had reached the interval goal, we found higher response rate compared with intervals where the goal was not reached. This is reflected in both (B) faster trial-wise reaction time for correct trials and (C) higher trial accuracy, though accuracy did not differ based on reward level. Error bars reflect standard errors. Asterisks denote the significance level of main effects. ***: *p* < 0.001

**Table 12.** Mixed Model Results for Correct Responses per Second Based on Challenge Level and Reward Level (Experiment 2)

| *Predictors* | Response Rate |  |  |
| --- | --- | --- | --- |
|  | *Estimates* | *SE* | *p-value* |
| Hard - Easy | 0.06 | 0.00 | < **0.001\*\*\*** |
| High Reward – Low Reward | 0.01 | 0.00 | < **0.001\*\*\*** |
| Interval Goal Completion | 0.41 | 0.01 | < **0.001\*\*\*** |
| Average Congruency | 0.00 | 0.00 | **0.043\*** |
| Scaled Interval Session Num | 0.02 | 0.00 | < **0.001\*\*\*** |
| Scaled Interval Block Num | 0.00 | 0.00 | 0.232 |
| Challenge X Reward | -0.00 | 0.00 | 0.333 |

**Table 13.** Mixed Model Results for Reaction Time and Accuracy based on Challenge and Reward Level (Experiment 2)

| *Predictors* | Response Time<br>(for accurate trials) |  |  | Accuracy |  |  |
| --- | --- | --- | --- | --- | --- | --- |
|  | *Estimates* | *SE* | *p-value* | *Log Odds Ratios* | *SE* | *p-value* |
| Hard - Easy | -31.07 | 3.94 | < **0.001\*\*\*** | 0.50 | 0.03 | < **0.001\*\*\*** |
| High Reward – Low Reward | -4.92 | 1.54 | **0.001\*\*\*** | 0.01 | 0.02 | 0.693 |
| Trial Goal Completion | -11.96 | 6.60 | 0.070 | 1.32 | 0.07 | < **0.001\*\*\*** |
| Interval Goal Completion | -168.01 | 15.38 | < **0.001\*\*\*** | 1.82 | 0.07 | < **0.001\*\*\*** |
| Scaled Trial Interval Num | -242.69 | 8.24 | < **0.001\*\*\*** | -0.14 | 0.11 | 0.202 |
| Dummy Trial Num | 265.66 | 10.66 | < **0.001\*\*\*** | -0.63 | 0.15 | < **0.001\*\*\*** |
| Scaled Trial Session Num | -32.88 | 8.51 | < **0.001\*\*\*** | -0.25 | 0.12 | **0.036\*** |
| Scaled Interval Session Num | 21.44 | 8.27 | **0.010\*\*** | 0.26 | 0.11 | **0.023\*** |
| Scaled Interval Block Num | -2.24 | 1.28 | 0.081 | 0.00 | 0.02 | 0.783 |
| Challenge X Reward | 1.49 | 1.52 | 0.326 | -0.02 | 0.02 | 0.425 |
| Challenge X Trial Goal Completion | 0.03 | 3.32 | 0.992 | -0.03 | 0.05 | 0.531 |
| Reward X Trial Goal Completion | -0.69 | 3.17 | 0.827 | 0.05 | 0.05 | 0.320 |
| Scaled Trial Interval Num X Dummy Trial Num | 251.98 | 8.23 | < **0.001\*\*\*** | -0.98 | 0.11 | <**0.001\*\*\*** |

#### Effects of goal completion and proximity on performance

As in Experiment 1, we examined effects of goal completion and proximity, focusing only on intervals in which the minimal goal was met. Similar to our findings before, participants were faster to complete trials correctly (*F*(1, 157) = 6.61, *p* = 0.010) and more accurate (*X^2^*(1, N = 38) = 233.12, *p* < 0.001) after they had reached the minimum goal (Table 14, Figure 8).

**Figure 8.**
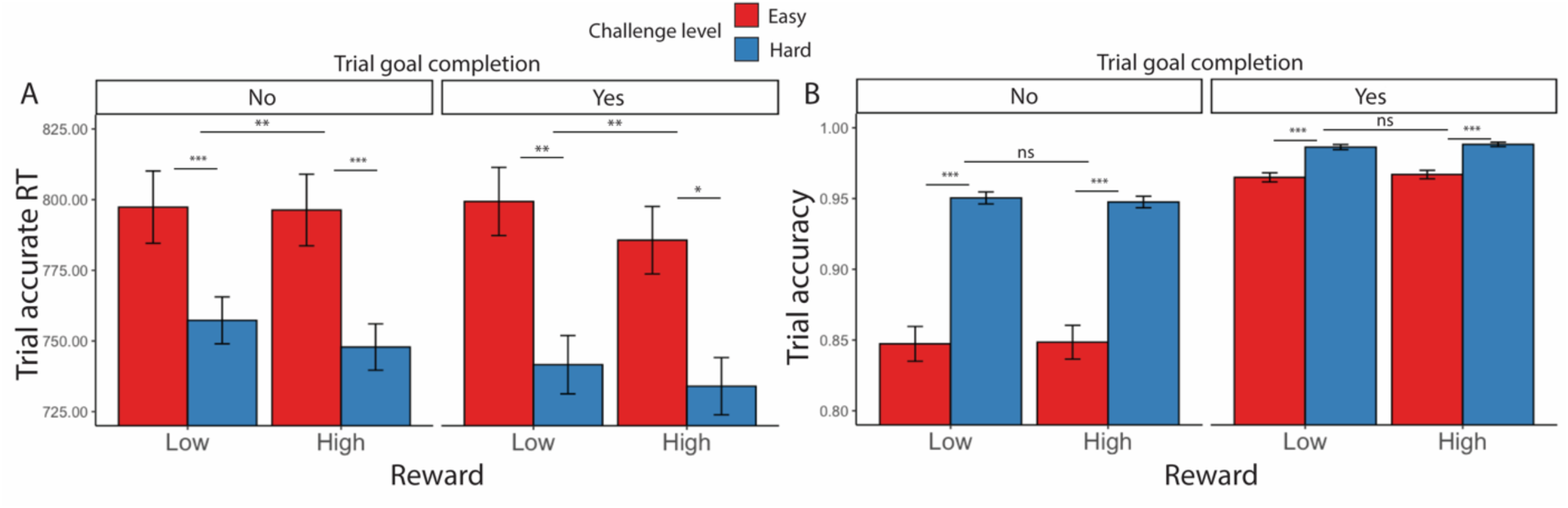
Goal completion effects (Experiment 2). Similar to Experiment 1, participants are (A) overall faster in completing correct trials and (B) more accurate after they had reached the interval goal. Error bars reflect standard errors. Asterisks denote the significance level of main effects. ***: *p* < 0.001; **: *p* < 0.01; *: *p* < 0.05

**Table 14.** Mixed Model Results for Reaction Time and Accuracy based on Challenge and Reward Level (Experiment 2, goal-reached intervals only)

| *Predictors* | Response Time<br>(for accurate trials) |  |  | Accuracy |  |  |
| --- | --- | --- | --- | --- | --- | --- |
|  | *Estimates* | *SE* | *p-value* | *Log Odds Ratios* | *SE* | *p-value* |
| Hard - Easy | -27.35 | 3.08 | < **0.001\*\*\*** | 0.52 | 0.04 | < **0.001\*\*\*** |
| High Reward – Low Reward | -5.40 | 1.44 | < **0.001\*\*\*** | 0.01 | 0.03 | 0.607 |
| Trial Goal Completion | -16.16 | 6.28 | **0.010\*\*** | 1.37 | 0.09 | <**0.001\*\*\*** |
| Scaled Trial Interval Num | -226.65 | 7.88 | < **0.001\*\*\*** | -0.11 | 0.14 | 0.436 |
| Dummy Trial Num | 250.65 | 10.20 | < **0.001\*\*\*** | -0.77 | 0.19 | <**0.001\*\*\*** |
| Scaled Trial Session Num | -20.16 | 8.43 | **0.017\*** | -0.18 | 0.14 | 0.200 |
| Scaled Interval Session Num | 7.86 | 8.32 | 0.345 | 0.20 | 0.14 | 0.162 |
| Scaled Interval Block Num | -2.36 | 1.21 | 0.051 | 0.02 | 0.02 | 0.244 |
| Challenge X Reward | -0.32 | 1.43 | 0.824 | -0.03 | 0.03 | 0.208 |
| Challenge X Trial Goal Completion | -1.91 | 2.97 | 0.520 | -0.08 | 0.05 | 0.132 |
| Reward X Trial Goal Completion | -0.06 | 2.86 | 0.982 | 0.04 | 0.05 | 0.407 |
| Scaled Trial Interval Num X Dummy Trial Num | 237.99 | 7.88 | < **0.001\*\*\*** | -1.15 | 0.14 | <**0.001\*\*\*** |

We found similar goal proximity effects as in Experiment 1 (Table 15). As seen in Figure 9, we again found that participants initially sped up after starting the interval for approximately 2 trials, then gradually slowed as they neared and surpassed the goal. After excluding the first two trials and the last trial in each interval, we again found that participants were slower in completing correct trials near the goal (*F*(1, 124.9) = 113.82, *p* < 0.001). We also found that accuracy decreased across conditions as participants approached the goal (*X^2^*(1, N = 78) = 5.65, *p* = 0.012), something that we only found reliably for more challenging intervals in Experiment 1. As in Experiment 1, we also found an interaction between goal proximity and challenge level on trial accuracy, such that this decrease in accuracy was steeper for more challenging intervals (*X^2^*(1, N = 78) = 4.85, *p* = 0.028), though again we cannot rule out explanations for this interaction related to within-interval fatigue.

**Figure 9.**
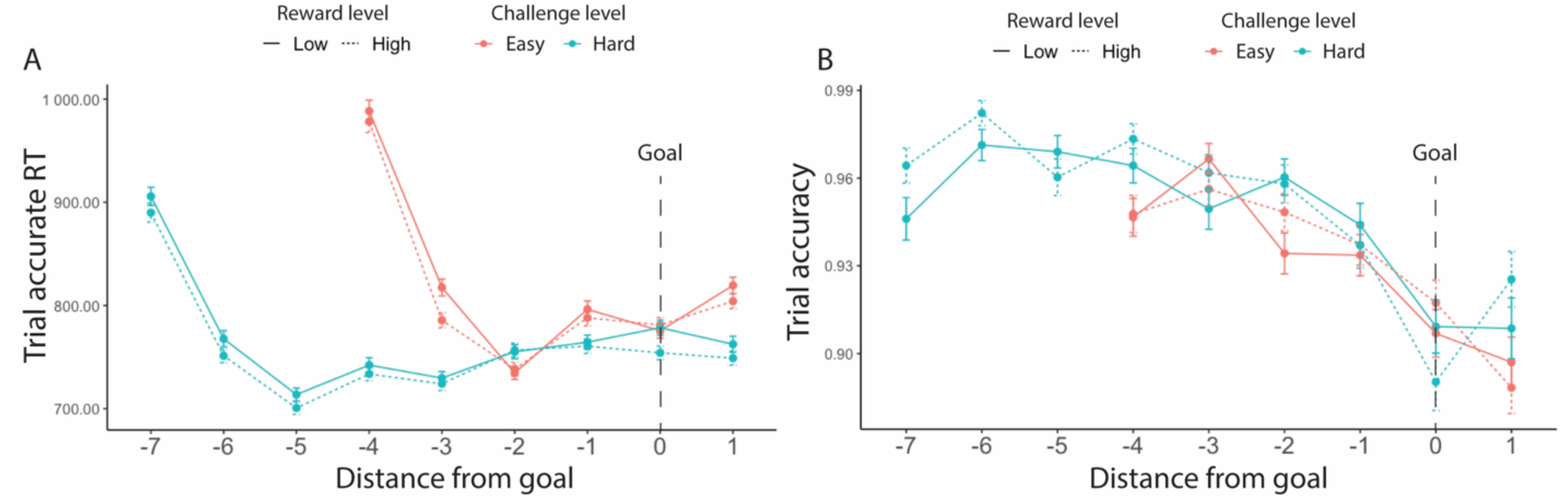
Goal proximity effects in reaction time (Experiment 2). We found similar results to Experiment 1: Overall, participants were slower when they approached the goal. Their accuracy also decreased. We again saw that participants initially sped up in performing accurate trials after starting the interval but then slowed down (at point -2 for Easy and -5 for Hard). This decreasing of speed extended until they had surpassed the goal. Error bars reflect standard errors.

**Table 15.** Mixed Model Results for Goal Proximity Effect for Reaction Time and Accuracy (Experiment 2)

| *Predictors* | Response Time (for accurate trials) |  |  | Accuracy |  |  |
| --- | --- | --- | --- | --- | --- | --- |
|  | *Estimates* | *SE* | *p-value* | *Log Odds Ratios* | *SE* | *p-value* |
| Goal Distance | -20.38 | 1.91 | < **0.001\*\*\*** | 0.13 | 0.05 | **0.012\*** |
| Hard - Easy | -15.10 | 2.31 | < **0.001\*\*\*** | 0.25 | 0.04 | < **0.001\*\*\*** |
| High Reward – Low Reward | -4.92 | 1.56 | **0.002\*\*** | 0.02 | 0.03 | 0.622 |
| Scaled Trial Session Num | -18.41 | 9.17 | **0.045\*** | -0.69 | 0.16 | < **0.001\*\*\*** |
| Scaled Interval Session Num | 6.94 | 9.11 | 0.446 | 0.69 | 0.16 | < **0.001\*\*\*** |
| Scaled Interval Block Num | -1.69 | 1.32 | 0.201 | 0.03 | 0.03 | 0.242 |
| Goal Distance X Challenge | 2.73 | 1.80 | 0.129 | 0.08 | 0.04 | **0.028\*** |
| Goal Distance X Reward | -2.05 | 1.76 | 0.244 | 0.03 | 0.03 | 0.351 |
| Challenge X Reward | 0.70 | 1.54 | 0.650 | 0.03 | 0.03 | 0.306 |

Together, we found that higher reward encouraged participants to perform better and persist more with their cognitive effort, as was the case for more challenging intervals in this experiment and the previous one. Within an interval, we saw similar patterns as Experiment 1, such that performance improved after the goal was reached.

#### Effects of reward and challenge level on affective experiences

Similar to Experiment 1, we found that participants reported feeling better (*F*(1, 100.6) = 98.86, *p* < 0.001) and less stressed (*F*(1, 93.08) = 33.80, *p* < 0.001) after reaching the goal (Table 16). Focusing on intervals where the minimum goal had been reached, participants reported higher stress (*F*(1, 92.5) = 44.69, *p* < 0.001) and less positive affect (*F*(1, 88.7) = 44.52, *p* < .001) for hard relative to easy intervals (Figure 10, Table 17). When examining the effect of reward over and above these challenge level effects, we found that high reward relative to low reward intervals led participants to report feeling more positive (*F*(1, 118.8) = 6.20, *p* = 0.014) but *also* more stressed (*F*(1, 129.5) = 18.84, *p* < .001). We also found an interaction between reward and challenge level in predicting stress ratings, whereby higher reward enhanced the stress and positive affect induced by more challenging intervals (controlling for the influence of task performance; *F*(1, 1869.43) = 4.88, *p* = 0.040; Table 17).

**Figure 10.**
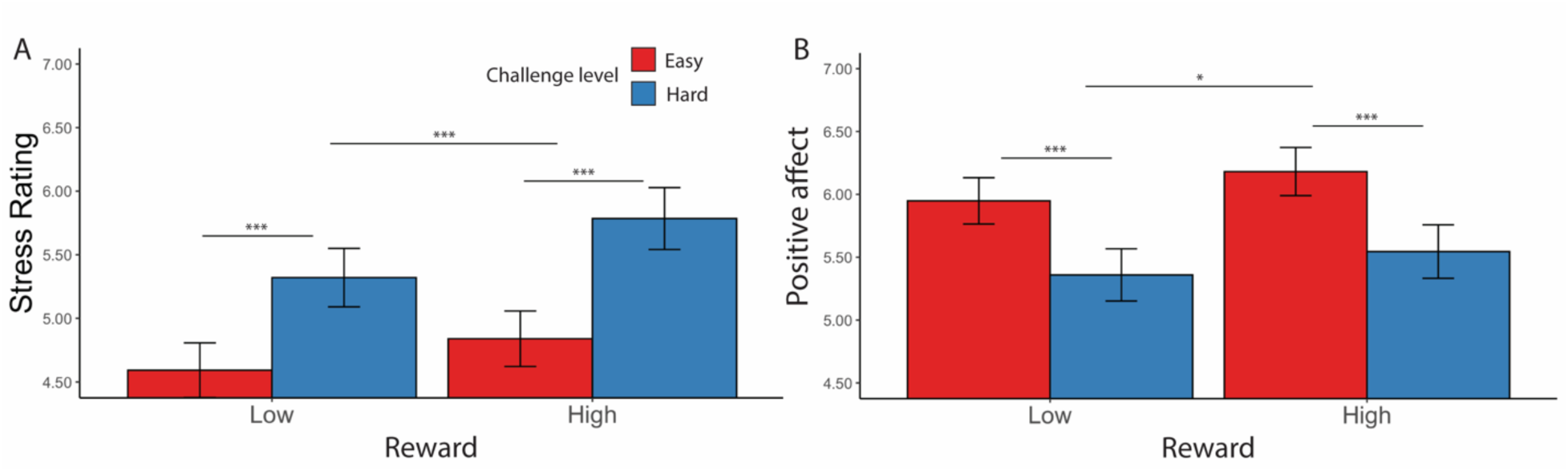
Affective Ratings based on challenge level and reward level, for goal completed intervals only. (A) Stress Ratings: Participants reported feeling more stress after completing hard intervals and intervals with higher potential reward. (B) Positive Affect: Participants reported feeling better after completing intervals with higher potential reward, but there was no difference in positive affect with respect to challenge level. Error bars reflect standard errors. Asterisks denote the significance level of main effects. ***: *p* < 0.001; *: *p* < 0.05

**Table 16.** Mixed Model Results for Affective Ratings Based on Challenge Level, Reward, and Goal Completion (Experiment 2)

| Predictors | Stress |  |  | Positive Affect |  |  |
| --- | --- | --- | --- | --- | --- | --- |
|  | Estimates | SE | p-value | Estimates | SE | p-value |
| Hard - Easy | 0.38 | 0.15 | **0.015\*** | 0.07 | 0.16 | 0.657 |
| High Reward – Low Reward | 0.08 | 0.14 | 0.550 | -0.20 | 0.15 | 0.185 |
| Interval Goal Completion | -1.56 | 0.27 | **< 0.001\*\*\*** | 3.16 | 0.32 | **< 0.001\*\*\*** |
| Average Congruency | -0.02 | 0.03 | 0.646 | 0.05 | 0.03 | 0.180 |
| Scaled Interval Session Num | -0.23 | 0.03 | **< 0.001\*\*\*** | -0.12 | 0.03 | **< 0.001\*\*\*** |
| Scaled Interval Block Num | 0.04 | 0.03 | 0.202 | -0.07 | 0.03 | **0.046\*** |
| Challenge X Reward | 0.01 | 0.14 | 0.958 | 0.09 | 0.15 | 0.537 |
| Challenge X Interval Goal Completion | -0.13 | 0.15 | 0.393 | -0.13 | 0.16 | 0.424 |
| Reward X Interval Goal Completion | 0.05 | 0.14 | 0.733 | 0.36 | 0.16 | **0.022\*** |

**Table 17.** Mixed Model Results for Affective Ratings Based on Challenge and Reward Level, Controlling for Performance (Goal Completed Intervals Only, Experiment 2)

| Predictors | Stress |  |  | Positive Affect |  |  |
| --- | --- | --- | --- | --- | --- | --- |
|  | Estimates | SE | p-value | Estimates | SE | p-value |
| Response Rate | -0.88 | 0.10 | **< 0.001\*\*\*** | 1.32 | 0.14 | **< 0.001\*\*\*** |
| Hard - Easy | 0.44 | 0.07 | **< 0.001\*\*\*** | -0.35 | 0.05 | **< 0.001\*\*\*** |
| High Reward – Low Reward | 0.18 | 0.04 | **< 0.001\*\*\*** | 0.10 | 0.04 | **0.013\*** |
| Average Congruency | 0.01 | 0.03 | 0.867 | 0.01 | 0.03 | 0.804 |
| Scaled Interval Session Num | -0.14 | 0.03 | **< 0.001\*\*\*** | -0.21 | 0.03 | **< 0.001\*\*\*** |
| Scaled Interval Block Num | 0.02 | 0.03 | 0.518 | -0.06 | 0.03 | **0.039\*** |
| Response Rate X Challenge | -0.09 | 0.06 | 0.127 | 0.17 | 0.05 | **0.001\*\*\*** |
| Response Rate X Reward | -0.02 | 0.05 | 0.694 | -0.00 | 0.05 | 0.978 |
| Challenge X Reward | 0.08 | 0.04 | **0.040\*** | -0.01 | 0.04 | 0.693 |

**Table 18.** Mixed Model Results for Affective Ratings Based on Average Reaction Time (Goal Completed Intervals Only, Experiment 2)

| *Predictors* | Stress |  |  | Positive Affect |  |  |
| --- | --- | --- | --- | --- | --- | --- |
|  | *Estimates* | *SE* | *p-value* | *Estimates* | *SE* | *p-value* |
| Mean Accurate RT | 0.52 | 0.10 | < **0.001\*\*\*** | -0.76 | 0.12 | < **0.001\*\*\*** |
| Hard - Easy | 0.34 | 0.06 | < **0.001\*\*\*** | -0.15 | 0.05 | **0.002\*\*** |
| High Reward – Low Reward | 0.19 | 0.04 | < **0.001\*\*\*** | 0.15 | 0.05 | **0.002\*\*** |
| Average Congruency | -0.00 | 0.04 | 0.998 | 0.03 | 0.04 | 0.445 |
| Scaled Interval Session Num | -0.17 | 0.04 | < **0.001\*\*\*** | -0.19 | 0.04 | < **0.001\*\*\*** |
| Scaled Interval Block Num | 0.02 | 0.03 | 0.648 | -0.06 | 0.04 | 0.067 |
| Mean Accurate RT X Challenge | 0.09 | 0.06 | 0.146 | -0.01 | 0.06 | 0.923 |
| Mean Accurate RT X Reward | 0.06 | 0.05 | 0.268 | 0.01 | 0.06 | 0.810 |
| Challenge X Reward | 0.10 | 0.04 | **0.012\*** | -0.03 | 0.04 | 0.490 |

For goal-completed intervals, we also replicated Experiment 1’s finding that participants reported better affective experiences (i.e., less stress and greater positive affect) with better performance, as measured by response rate (less stress: *F*(1, 89.3) = 79.89, *p* < 0.001; greater positive affect: *F*(1, 81.88) = 91.55, *p* < 0.001), trial-wise accuracy (less stress: *F*(1, 90.50) = 91.29, *p* < 0.001; greater positive affect: *F*(1, 73.78) = 88.98, *p* < 0.001), and reaction time of correct trials (less stress: *F*(1,86.72) = 24.81, *p* < 0.001; greater positive affect: *F*(1, 84.68) = 41.64, *p* < 0.001) (Table 17-19, Figure 11). We again found that the challenge-induced affective experience (i.e., higher stress, less positive affect) was reduced as participants performed better, once again relating specifically to accuracy rather than reaction time (*F*(1, 1503.62) = 5.11, *p* = 0.024, Table 19).

**Figure 11.**
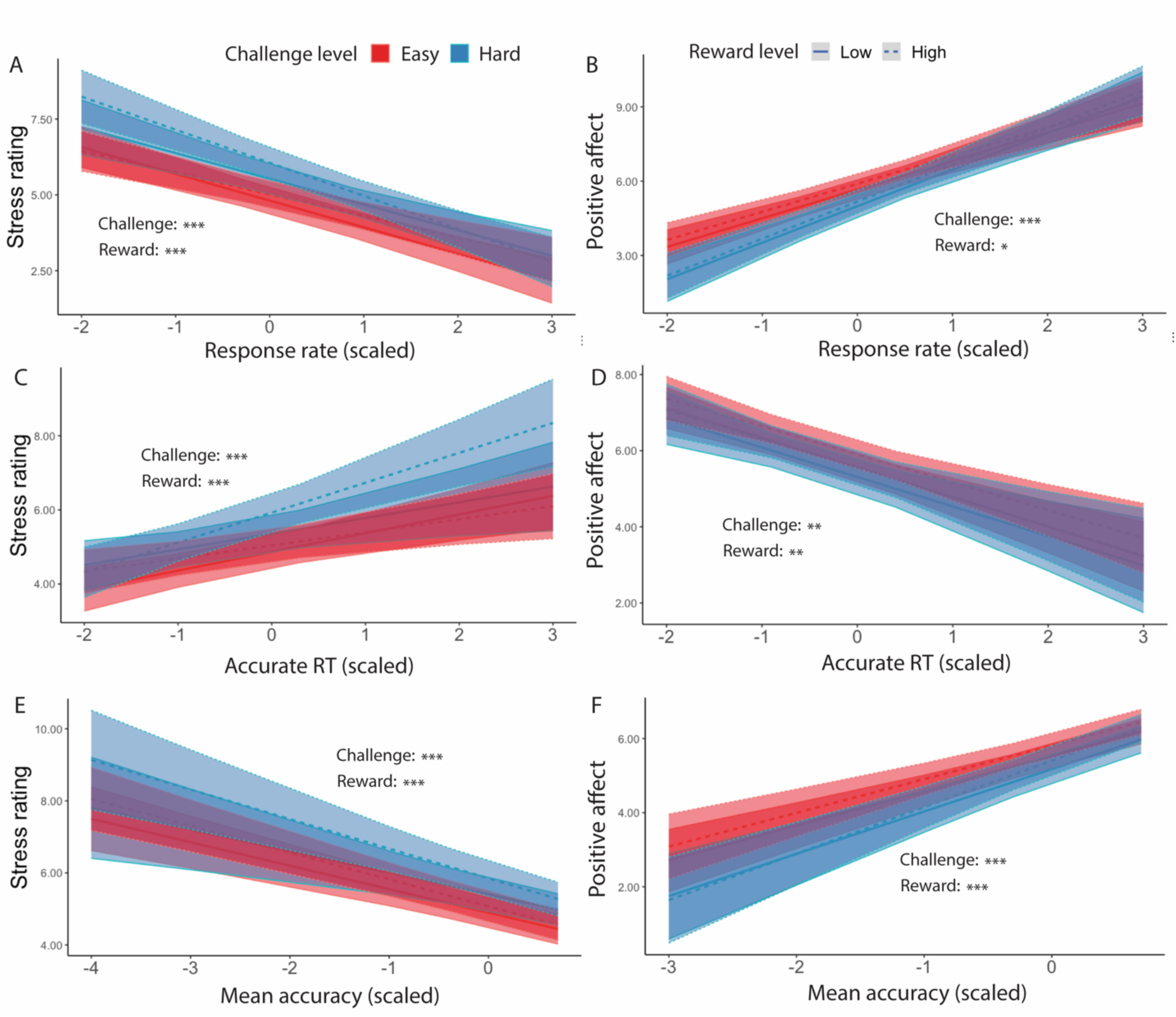
Affective Ratings predicted by performance, for goal-completed intervals only (Experiment 2). We found that in intervals where the goal had been reached, participants reported to have felt less stressed when (A) performed more correct trials per interval, (C) faster reaction time, and (E) higher accuracy. They also reported feeling better overall when (B) performed more correct trials per interval, (D) faster reaction time, and (F) higher accuracy. While we found that challenge level interacted with response rate and accuracy in predicting positive affect, there was no interaction of reward with performance. Error bars reflect standard errors. Asterisks denote the significance level of main effects. ***: *p* < 0.001; **: *p* < 0.01; *: p < 0.05

**Table 19.**
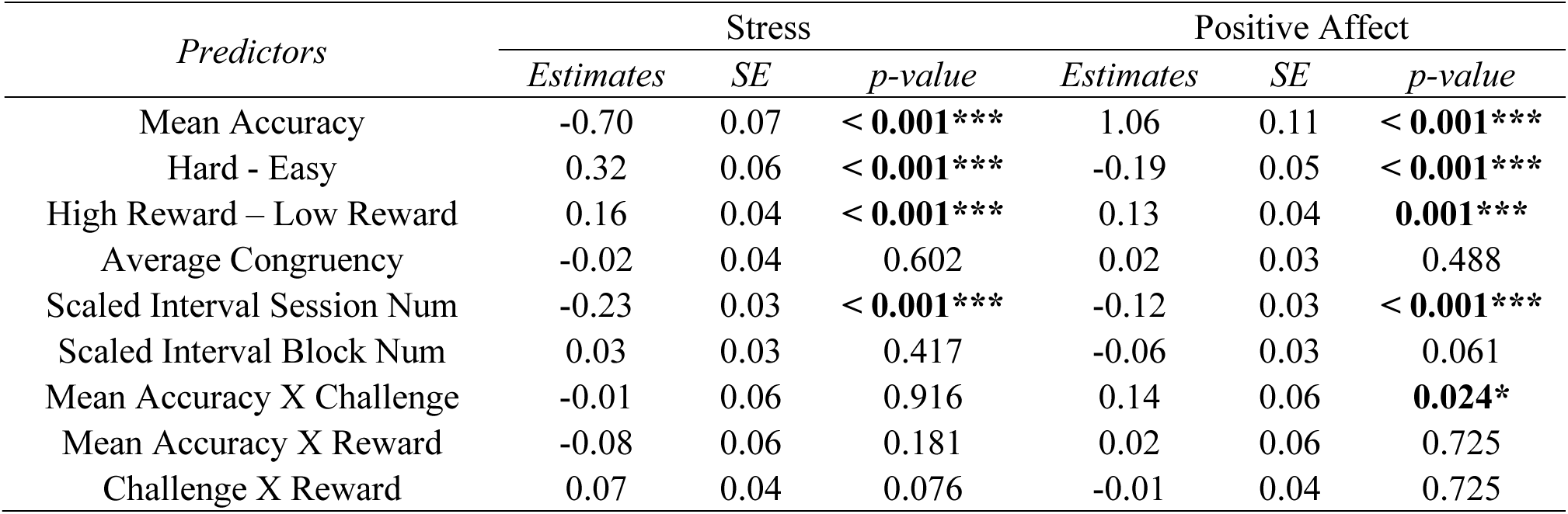
Mixed Model Results for Affective Ratings Based on Average Accuracy (Goal Completed Intervals Only, Experiment 2)

### Experiment 2 Discussion

By varying reward level in addition to challenge level, we were able to show that both variables motivated better performance, with participants completing more correct trials per interval with higher levels of expected reward and challenge, respectively. Within intervals, the effect of goal completion on increasing trial-wise accuracy and speeding up reaction time largely replicated that in Experiment 1, with higher rewards also exerting an additive rather than interactive influence on those RTs, but not accuracy, over and above the influence of challenge level. These findings suggest that both higher challenge and greater reward may similarly enhance selective attention, while reward could also reduce response caution (Grahek et al., 2024; Leng et al., 2021). We largely replicated the effects of goal proximity here, such that participants performed the task more slowly as they approached the goal, and in this experiment additionally found that they were less accurate as they approached the goal. We found that higher levels of expected reward, like higher levels of expected challenge level, induced greater stress, while also contributing to greater positive affect when the participant met their goal. We again found that better performance, particularly reflected in higher accuracy, predicted feeling more positive and less stressed.

## Discussion

We engage in effortful control to reach goals constantly. The amount of control invested depends on motivation to achieve the goal, which is determined by factors such as performance incentives and task demands (Botvinick & Braver, 2015). The current work investigated the role of expected challenge and reward level on effort exertion and persistence in a cognitively demanding task, as well the affective experience associated with performing the task. Overall, participants were motivated to allocate more cognitive effort, and prolong persistence of effort, when faced with more challenging goals and higher potential rewards, resulting in better performance on the task. Participants experienced greater stress when performing under greater challenge and/or higher stakes, but these experiences were accompanied with greater positive affect when the stakes were high, and when the participant performed well during that interval (which also mitigated stress levels).

### Task performance varies with challenge, reward, and goal proximity

The relationship we found between challenge level and task performance support a central prediction of the goal setting theory, that more challenging goals lead people to work harder. As Locke et al. (1981) found, more challenging goals produce better performance because people exert more effort to complete it. This is reflected in our study, as both of our experiments found that participants completed more correct trials per interval in more challenging intervals, even though individual trials were rewarded *equally* across easy and hard conditions. Results from our Experiment 2 additionally pointed to monetary incentives as a motivator of performance and goal attainment, which is consistent with past work on goal setting theory and research on the incentivization of cognitive control (Botvinick & Braver, 2015; Wright, 1992).

The goal threshold and timed-interval design of our study also allowed us to examine how effort persistence is influenced by challenging goals. Because participants are free to choose how many trials they want to perform within these fixed time intervals, they could have chosen to stop exerting effort once the goal had been reached, or even prior to reaching the goal if they found it too taxing or impossible to reach. However, our results show that participants not only continued to perform more trials after reaching the goal, but they were also faster to complete trials correctly and more likely to respond correctly after the goal had been reached. Therefore, our results support that adequate expected challenge enhances effort exertion and prolongs cognitive effort persistence.

Across both experiments, we found that participants slowed down as they neared the goal, indicating more cautious behavior as they approached goal completion. Interestingly, we found that participants initially sped up in performing accurate trials after starting the interval but then slowed down. This dynamic may reflect the change in the perceived likelihood of completing the goal, such that participants begin each interval with an initial under-estimated chance of completion, leading to initial speeding. In contrast, the follow-up slowing could reflect the reversal of speeding associated with over-estimated chance of completion as they approach the goal. Such distinct patterns of speeding and slowing over the course of the interval, could also reflect dynamic reconfigurations of control across information processing (e.g., enhancing the accumulation of incoming evidence) and response thresholds (Grahek et al., 2024; Ritz et al., 2022). Indeed, recent work suggests that participants adjust both their evidence accumulation rate and their response threshold as they approach a goal (albeit over longer timescales than in the current study) (Devine et al., 2024). It will be valuable to test whether the current effects result from similar or distinct dynamics (e.g., differential changes in drift rate vs. threshold), the extent these parameters vary with reward and challenge level, and whether these can be collectively accommodated by normative models of control allocation (e.g., based on optimization of effort-discounted reward rate) (Leng et al., 2021; Prater Fahey et al., 2023).

### Affective experiences vary with challenge, reward, goal attainment, and performance

Goal attainment has been found to be related to satisfaction and well-being (Parker et al., 2009). In our study, goal attainment promoted greater positive affect, as did the prospect of gaining higher monetary reward. We also found that failure to attain a goal led to higher stress. Interestingly, our results indicate that the effects of goal attainment on affective experience are additionally impacted by challenge level, as harder challenge led participants to feel more stressed and less positive affect, despite having completed the task goal. Higher reward, on the other hand, led to higher stress and greater positive affect, which might appear paradoxical. This could reflect dual appraisals of the rewards at stake in terms of their consummatory value as well as the potential opportunity cost for completing fewer trials than one is able (Shenhav et al., 2014). Therefore, more potential reward would cause more stress during task performance, but also lead to greater satisfaction when received. Whether participants are framing these stakes as potential losses, and if this framing has any effect on behavior, is worth exploring in future studies (Prater Fahey et al., 2023).

The effects of goal attainment on affective experience were influenced by how much effort the participant had put into the task, as measured by response rate, average reaction time, and average accuracy. Across both experiments, participants felt more positive and less stressed when they had exerted greater and more prolonged cognitive effort to achieve better performance and had reached their goals. Interestingly, in Experiment 1, we also found that expected challenge level interacted with performance in predicting affective ratings – performing well had a more salutary effect on affect (greater increases in positive affect and decreases in stress) for hard than easy intervals. This suggests that when participants view a task as more challenging, they place more value in the effort they exerted (Inzlicht et al., 2018). Notably, this interaction effect was only found with task accuracy and not reaction time, suggesting that affective experience might be selectively attached to how accurate the performance was rather than simply how fast they are performing. Taken together, our results suggest that participants’ experiences of stress and positive affect not only take into account the end result, but also the process and intrinsic motivation that led there.

The findings that higher challenge and reward levels both led to greater levels of stress are also consistent with the affective patterns seen in the phenomenon of choking under pressure, when facing challenging and high stakes tasks could lead to increased stress and anxiety (Baumeister & Showers, 1986). In contrast to the unexpected decrease in performance as seen in the choking phenomenon, our task induced a level of pressure that instead motivated performance and cognitive control engagement compared with a low pressure condition. This could be that the amount of pressure faced by participants did not significantly alter the amount of attention, as suggested by the Distraction theory (Eysenck et al., 2007; Lee & Grafton, 2015) or Explicit Monitoring theory (Lewis & Linder, 1997), or level of arousal, as suggested by the Over Motivation theory (Ariely et al., 2009; Easterbrook, 1959), to the extent that it is crippling to task performance. Future studies could potentially disentangle these possibilities by modifying the task structure (e.g., setting the threshold higher to induce greater pressure) and evaluating the performance and affective outcomes.

### Limitations and Future Directions

Our task relies on self-report measures of affective experiences, measured repeatedly throughout the experimental session, at the end of each interval. In spite of its benefits, this approach risks eliciting demand characteristics related to inferences of what affective ratings we as experimenters believe to be relevant to the different experimental conditions. Conversely, giving these ratings may diminish participants’ affective experiences (Kassam & Mendes, 2013; Torre & Lieberman, 2018). Future studies could aim to incorporate physiological measures of acute stress (e.g., skin conductance, EKG) for a more well-rounded perspective.

Our goals were also uniform across participants, always requiring 5 correct responses for easy intervals and 8 correct responses for hard intervals. Though we chose these thresholds based on their ability to generate intervals that were generally achievable for the majority of participants, but differing in the effort required to meet that challenge, the absence of individualized goals meant that the level of challenge and achievability likely differed across participants. Future studies should aim to calibrate these thresholds to a given participant. In addition, selection of other threshold values could lead to difference in performance and affective experiences that could be of interest for future studies (e.g., requiring 10 correct responses instead of 8 likely leads to greater number of unreachable intervals). Additionally, because of the overall high goal completion rates in our data, we had insufficient intervals to tease apart the behavioral strategies that resulted in participants failing to reach their goal. For instance, it could be that failures to reach a goal reflected intervals where the participant was trying to meet the goal but felt unable to meet the challenge, for example due to capacity limitations or fatigue. Alternatively, failure to reach the goal could reflect slacking rather than an inability to achieve the goal, especially when these failures occurred for easy intervals, where participants generally performed near ceiling in terms of goal attainment. These two types of goal failure can yield different patterns of performance and affective experience, and future work should aim to disentangle these.

In conclusion, our study provides evidence that more challenging goals lead people to work harder but to also experience greater stress doing so, and that promising greater rewards lead to an enhancement rather than diminution of both of these effects. The wide application of goal setting theory and the influence of challenge level in fields such as education and industry, in motivating students and workers to perform their best, prompts more extensive studies on this topic. Focusing especially on how acute stress, resulting from the goal, task, and potential reward, plays a role in effort motivation is important not only in promoting performance, but also in ensuring the mental health and well-being of those who are completing the tasks.

## Data availability statement

Data and analysis code are available online (https://github.com/yzhangl/TSSS_Materials.git).

## Author contributions

This study was conceived by Y.Z. and A.S., the task was programmed by Y.Z. and X.L., and the data were collected and analyzed by Y.Z., under the supervision of X.L. and A.S. Y.Z. drafted the manuscript and all authors contributed to revisions.

## Acknowledgements and funding information

This research was supported by an NIH T32-MH115895 (X.L.) and an NSF CAREER Award 2046111 (A.S.). We are grateful to Mahalia Prater Fahey, Debbie Yee, and Amanda Arulpragasam for helpful discussions and feedback.

## Ethics approval

Approval was obtained from Brown University’s Institutional Review Board. The procedures used in this study adhere to the tenets of the Declaration of Helsinki.

## Conflicts of interest/Competing interests

The authors have no relevant financial or non-financial interests to disclose.

## Footnotes

1 We use the phrase “challenge level” to distinguish from “task difficulty” as noted in Locke et al (1981). The authors had coined the term difficulty to refer to the difficulty of the task itself (e.g., writing a novel is much harder than writing a birthday card). On the other hand, challenge level usually refers to attaining a specific standard of proficiency within a time limit (e.g., completing 20 versus 5 simple arithmetic questions within 10 minutes has distinct challenge level), which is what we manipulated in our tasks.

## Notes

https://github.com/yzhangl/TSSS_Materials

### Competing Interest Statement

The authors have declared no competing interest.

### Summary of Updates

Methods and results updated, additional discussions added, removed supplement & incorporated information into main manuscript, author affiliations updated

